# Scalable Generation of Pseudo-Unipolar Sensory Neurons from Human Pluripotent Stem Cells

**DOI:** 10.1101/2022.03.24.485622

**Authors:** Tao Deng, Carlos A. Tristan, Claire Weber, Pei-Hsuan Chu, Seungmi Ryu, Vukasin M. Jovanovic, Pinar Ormanoglu, Prisca Twumasi, Jaehoon Shim, Selwyn Jayakar, Han-Xiong Bear Zhang, Sooyeon Jo, Ty C. Voss, Anton Simeonov, Bruce P. Bean, Clifford J. Woolf, Ilyas Singeç

## Abstract

Development of new non-addictive analgesics requires advanced strategies to differentiate human pluripotent stem cells (hPSCs) into relevant cell types amenable for translational research. Here, we developed a highly efficient and reproducible method that differentiates hPSCs into peptidergic and non-peptidergic nociceptors. By modulating specific cell signaling pathways, hPSCs were first converted into SOX10^+^ neural crest cells, followed by differentiation into sensory neurons with an *in vivo*-like pseudo-unipolar morphology. Detailed characterization confirmed that the hPSC-derived nociceptors displayed molecular and cellular features comparable to native dorsal root ganglion (DRG) neurons, and expressed high-threshold primary sensory neuron markers, transcription factors, neuropeptides, and over 150 ion channels and receptors, including critical pain-relevant drug targets (e.g., TRPV1, TAC1, CALCA, NA_V_1.7, NA_V_1.8). Moreover, after confirming robust functional activities and differential response to noxious stimuli and specific drugs, a robotic cell culture system was employed to produce large quantities of human sensory neurons, which can be used to develop nociceptor-selective analgesics.

## INTRODUCTION

Chronic pain affects over 30% of the world population representing a major unmet medical need^1,2^. Pain is a highly subjective experience and difficult to model during preclinical development. Traditional cellular models in pain research include primary rodent cells, heterologous expression of ion channels in non-neuronal cells (e.g. cancer cells, Xenopus oocytes), and in a few cases human postmortem DRG^3–5^. Animal models do not completely phenocopy human physiology and this may in part explain the poor efficacy or unexpected toxicity of many pain drug candidates in clinical trials^6,7^. Recent studies identified various species-specific differences at the molecular and cellular level in human and mouse DRG neurons, suggesting that the organizational principles of the sensory system in mouse and humans are substantially different^8–11^.

Nociceptors are specialized cell types of the peripheral nervous system (PNS) and critical for transmitting information on the presence of noxious stimuli from the body to the spinal cord and higher brain centers of the somatosensory system^12^. The cell bodies of nociceptors are localized to DRGs and one split axon of each nociceptor projects to the periphery as well as to the spinal cord where the first synaptic contact of the nociceptive pathway is established with dorsal horn neurons as postsynaptic partners. Unlike neurons in the central nervous system (CNS) such as multipolar cortical neurons with elaborate basal and apical dendritic trees, nociceptors are anatomically unique in that they exhibit a pseudo-unipolar morphology and lack dendrites. Nociceptors express a large number of ion channels, play a central role in inflammatory and neuropathic pain, and other chronic conditions due to congenital, metabolic, or iatrogenic causes (e.g. pain due to channelopathies, diabetic polyneuropathy, chemotherapy-induced polyneuropathy).

Self-renewing human PSCs lines, including embryonic stem cells (hESCs) and induced pluripotent stem cells (iPSCs) represent an attractive source for generating human sensory neurons^7,13–19^. Alternative transdifferentiation approaches are based on forced expression of transcription factors and lineage reprograming in mature cells to generate nociceptor-like cells^20,21^. Despite this progress, currently available methods are variable, difficult to scale-up, and generate neurons that display only some aspects of human nociceptor physiology. Here, we present an efficient and reproducible differentiation method for peptidergic and non-peptidergic nociceptors derived for human PSCs. These cells can be generated in large quantities, cryopreserved using a recently developed CEPT small molecule cocktail^22^, and utilized for translational studies and screens.

## Results

### Scalable production of human sensory neurons from hPSCs

All hPSCs used in this study were routinely cultured under feeder-free chemically-defined conditions using E8 medium and vitronectin as a coating substrate. To ensure stress-free cell expansion, cells were passaged using EDTA, and treated with the CEPT small molecule cocktail for 24 h, which promotes cell viability and fitness^22^. To differentiate neurons, hPSCs were plated at 1.5 x 10^5^ cells/cm^2^ and treated with A83-01 (TGF-β inhibitor) and CHIR98014, a new GSK-3β inhibitor that activates the WNT signaling pathway (**Fig. 1a**). Of note, when using the Hotspot kinase assay to test target specificity against 369 human kinases representing all major human kinase families^23^, we found that CHIR98014 was more potent than CHIR99021, although both compounds showed identical target selectivity profiles (**Extended Data Fig. 1a,b and Supplementary Table 1**). Introducing CHIR98014 as a superior WNT agonist is important because it efficiently activates the WNT signaling pathway at lower concentrations than CHIR99021, which can be cytotoxic in the absence of knockout serum replacement^24^. When hPSCs growing in adherent cultures were treated with A83-01 and CHIR98014 for 10 days, a mixture of SOX10^+^ and SOX10^−^ cells emerged. Importantly, SOX10^+^ cells were exclusively present in areas where cells had spontaneously aggregated and formed three-dimensional (3D) structures (**Extended Data Fig. 2a**). These spheres also contained cells expressing the transcription factor ISL1 and the neuronal marker TUJ1 (TUBB3) (**Extended Data Fig. 2a**). When spheres were manually transferred and attached to Geltrex-coated plates, cells migrated out of the spheres and developed typical neuronal morphologies (**Extended Data Fig. 2b**). Based on these observations, we decided to dissociate the differentiating cells at day 3 and plate them into ultra-low attachment plates (AggreWell plates) to promote sphere formation (**Fig. 1a and Extended Data Fig. 2c**). These structures, which we termed ‘nocispheres’, were then treated until day 14 with a combination of four factors including CHIR98014, A83-01, DBZ (γ-secretase inhibitor), and PD173074 (**Fig. 1a**). The Hotspot kinase assay revealed that the PD173074 compound potently and specifically inhibited the FGF receptors 1/2/3. In contrast, the more commonly used SU5402 blocked FGFR1/2/3 but showed even stronger off-target activity against the TrkB and TrKC neurotrophin receptors (**Extended Data Fig. 3 and Supplementary Table 1**). On day 14, nocispheres were dissociated into single cells and plated on Geltrex-coated dishes for further differentiation and maturation in a medium consisting of PD0332991 (CDK4/6 inhibitor) and a combination of neurotrophic factors (**Fig. 1a**). Over the course of 2 weeks, the number of SOX10^+^ precursor cells decreased, whereas the number of neuronal cells expressing BRN3A (POU4F1), a specific transcription factor expressed by nociceptors, increased strongly (**Fig. 1b-d**). Accordingly, the number of Ki-67^+^ proliferating cells decreased during the differentiation of SOX10^+^ neural crest into neurons (**Extended Data Fig. 4a,b**). Since large quantities of human neurons are needed for high-throughput screening projects, the cell differentiation protocol was automated by using the CompacT SelecT platform (**Extended Data Fig. 5a**), a robotic cell culture system that allows reproducible scale-up and biomanufacturing of human cells^25^. By culturing cells in three bar-coded large T175 flasks and with minimal manual intervention, over 150 million sphere-derived cells could be generated in 14 days in one experimental run (**Extended Data Fig. 5b**). Because the robotic platform can culture 90 flasks in parallel^25^, cell numbers can theoretically be increased 30-fold. At day 14, large quantities of cells can be cryopreserved using the CEPT cocktail or further differentiated until day 28 and used for the experiments described below.

**Fig. 1:**
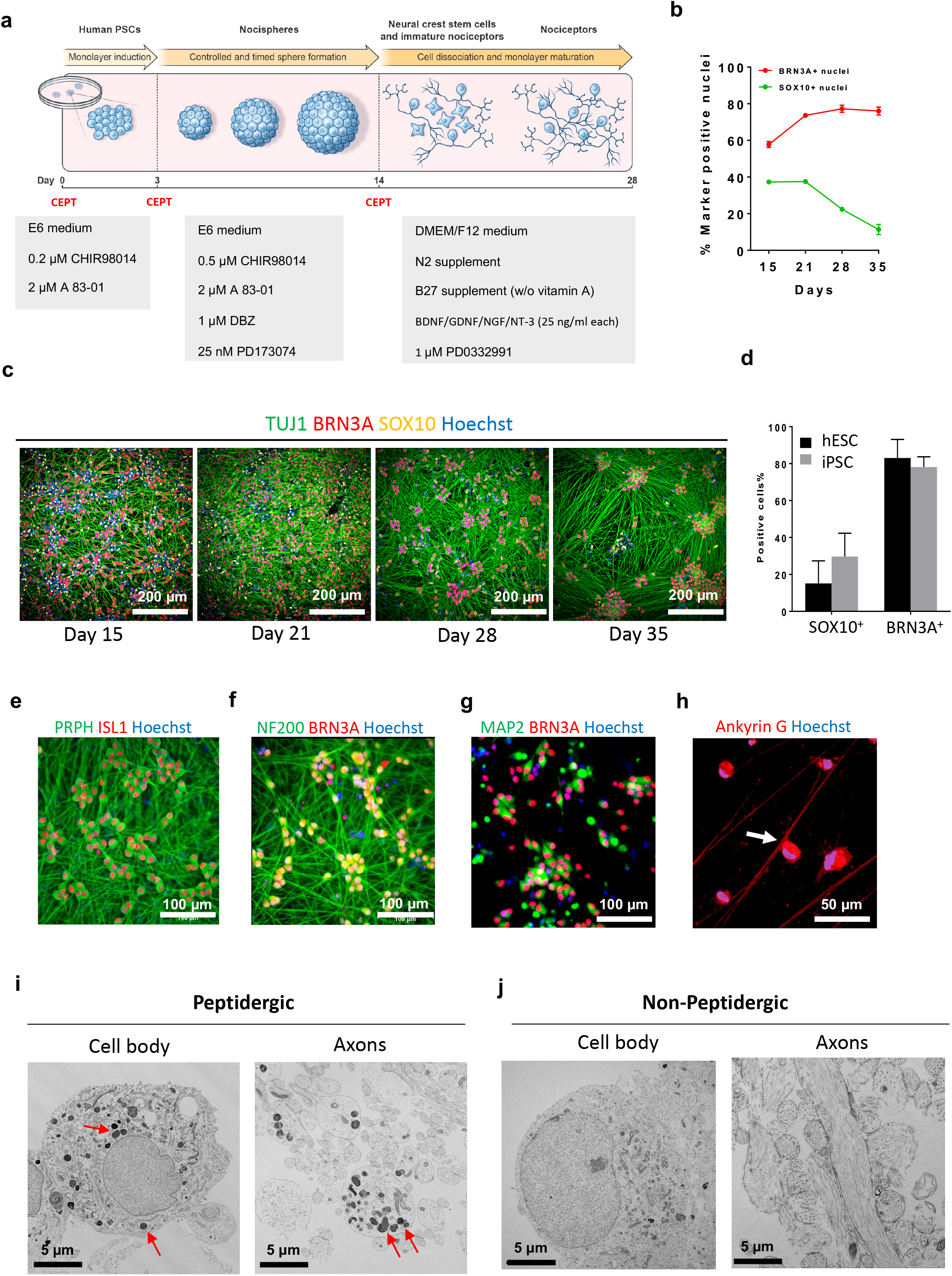
Directed differentiation of hPSC into pseudo-unipolar nociceptors. **a,** Schematic overview of nociceptor differentiation method. **b,** Quantitative analysis of cells expressing SOX10 and BRN3A during cell differentiation. Note that most cells express BRNA3A (a pan-sensory neuron marker) at day 28 and 35 (n = 3, mean ± s.d.) **c,** Representative images of differentiating cells at different timepoints immunolabeled for neural crest marker SOX10 and neuronal markers TUJ1 and BRNA3A. d, Quantification of cells expressing SOX10 or BRNA3A derived from hESCs (WA09) and iPSCs (LiPSC-GR1.1) showing consistency of the differentiation method (n = 3, mean ± s.d.). **e, f**, Representative images of nociceptor cultures at day 28 expressing the axonal markers PRPH (peripherin) and NF200, and transcription factors ISL1 and BRNA3A. **g,** Immunostaining for MAP2 and BRNA3A at day 28. Note that MAP2 immunoreactivity is confined to cell bodies. **h,** Immunostaining for Ankyrin-G and confocal microscopy showing axon splitting of nociceptors (see Extended Data Fig. 8 for confocal z-stack images). **i,** Electron microscopic images showing peptidergic nociceptors. Prominent large dense core vesicles (LDCVs; red arrows) are present in the cytoplasm of the soma and in axonal profiles sectioned in different planes. See other examples for peptidergic nociceptors in Extended Data Fig. 9. **j,** Ultrastructure of a non-peptidergic nociceptor soma and axonal profiles lacking LDCVs. See Extended Data Fig. 8 for additional examples.

### Morphological characterization of pseudo-unipolar sensory neurons

Immunocytochemical analysis demonstrated that hPSC-derived nociceptors expressed neuronal markers TUJ1, peripherin, neurofilament 200 (NEFH), Tau (MAPT), and transcription factors BRNA3A and ISL1 (**Fig. 1e,f and Extended Data Fig. 6a**). Like DRG neurons, *in vitro*-generated nociceptors were immunoreactive for the neurotransmitter glutamate and expressed glutamate vesicular glutamate transporter 1 (vGLUT1)(**Extended Data Fig. 6b,c**). In these cultures, gamma-aminobutyric acid (GABAergic) neurons were only found very sporadically and cells expressing tyrosine hydroxylase (TH), the rate-limiting enzyme for dopamine synthesis and a feature of low threshold C-fiber mechanoreceptors^26^, were not detected (**Extended Data Fig. 6d,e**). Neurons are polarized cells and MAP2 is a typical marker to label the somato-dendritic compartment of neurons in the CNS. Accordingly, immunostaining for MAP2 only labeled the cell bodies of nociceptors (**Fig. 1g and Extended Data 6f**). This is consistent with the pseudo-unipolar anatomy of DRG neurons, which develop a split axon but are devoid of dendrites. The peripheral part of the split axon extends to the periphery such as the skin as free nerve endings, whereas the central part of the same axon projects to the dorsal horn to establish synaptic contacts with spinal cord neurons. By using confocal microscopy and immunostaining for the axon initial segment marker ankyrin G^27^, we detected axon splitting in hPSC-derived sensory neurons (**Fig. 1h and Extended Data Fig. 7**). To further characterize these hPSC-derived sensory neurons, we performed an electron microscopic analysis using day 28 cultures (**Fig. 1i, j and Extended Data Figs. 8, 9**). At the ultrastructural level, two types of sensory neurons could be distinguished based on their morphology. The first presumably peptidergic cell type displayed irregularly shaped nuclei with nucleolemma invaginations, rich endoplasmic reticulum, and numerous large dense core vesicles (LDCVs) that were present in the soma as well as axons (**Fig. 1i and Extended Data Fig. 8a-c**). The second presumably non-peptidergic cell type exhibited round nuclei of various sizes, numerous mitochondria, and was devoid of LDCVs (**Fig. 1j and Extended Data Fig. 8d-f**). Notably, while large numbers of axonal profiles with and without LDCVs were detected in these cultures, dendritic structures were absent (**Fig. 1i, j and Extended Data Fig. 8**). Focusing on the axo-somatic region, we found ultrastructural evidence for an axon directly emanating from the neuronal cell body (**Extended Data Fig. 9a,b**). Synapses are distinct structures defined by pre- and postsynaptic elements and a synaptic cleft. In these pure sensory neuron cultures, synapses were not observed at the ultrastructural level, suggesting that postsynaptic neurons were not generated using our differentiation method (**Fig. 1i, j and Extended Data Figs. 8,9**). Together, these findings provide evidence for efficient and scalable differentiation of hPSCs into sensory neurons with typical *in vivo*-like anatomical and immunophenotypic features.

### Transcriptomic analysis of directed differentiation

Time-course transcriptomic profiling (RNA-seq) was performed to generate datasets that characterize the differentiation trajectory from pluripotent cells to neural crest and sensory neurons (**Fig. 2**). Principal component analysis (PCA) showed distinct molecular signatures of pluripotent cells (day 0) and of the differentiating cells harvested at day 4, 8, 12, 21, 28 and 56 (**Fig. 2a**). Next, unbiased comparison of transcriptomes was performed by using ARCHS4, which is a RNA-seq database including over 84,863 human samples, followed by gene enrichment analysis (ENRICHR)^28^. This approach revealed that “sensory neuron” was the top-category at day 28 and 56, supporting the notion that the correct cell type was generated during *in vitro* differentiation (**Fig. 2b**). Furthermore, gene expression changes were investigated at different timepoints by using a heatmap analysis (**Fig. 2c**). Specific genes representing distinct stages and cell types were expressed or downregulated in a time-dependent fashion, consistent with neural crest lineage specification. Hence, the directed differentiation led to a downregulation of pluripotency-associated genes *OCT4* (*POU5F1*) and *NANOG* and the induction of neural crest genes including *SOX10*, *SOX21*, *PAX3* (**Fig. 2c and Extended Data Fig. 10a,b**). Upon further differentiation, these genes were then downregulated, followed by strong induction of non-specific neuronal genes as well as nociceptor-specific genes including *RUNX1*, *BRN3A*, *ISL1*, *TRPV1* and other genes that indicated the presence of both peptidergic and non-peptidergic subpopulations (**Fig. 2c and Extended Data Fig. 10c-g**). *RUNX3* is important for the development of proprioceptive neurons^29^ and was absent throughout the entire differentiation period (**Fig. 2d**). The transcription factor *PRDM12* is important for specifying nociceptors and genetic mutations in this gene can lead to congenital insensitivity to pain^17,30^. Transcriptomic analysis showed that expression of *PRDM12* was strongly induced, reached maximum expression at day 12, and then sharply declined by day 21-28, consistent with a stage-specific developmental gene expression (**Fig. 2e**). The sensory neuron-specific actin-binding protein advillin (*AVIL*)^31^ was upregulated during the differentiation (**Fig. 2f**). Three genes with diverse functions in cell polarity and axonal growth and with important roles in neurological disorders and Wallerian degeneration (*DPYSL2*, *GAP43, NMNAT2*) were induced and strongly expressed across the differentiation period (**Fig. 2g-i**). Lastly, a time-dependent regulation of several neurotrophin receptors (*NTRK1*, *NTRK2*, *NTRK3*, *NGFR*) and three receptors for glial cell-derived neurotrophic factor (*GFRA1*, *GFRA2*, *GFRA3*) was confirmed (**Fig. 2j-m and Extended Data Fig. 10h-j**), consistent with sensory neuron development *in vivo*^32,33^.

**Fig. 2:**
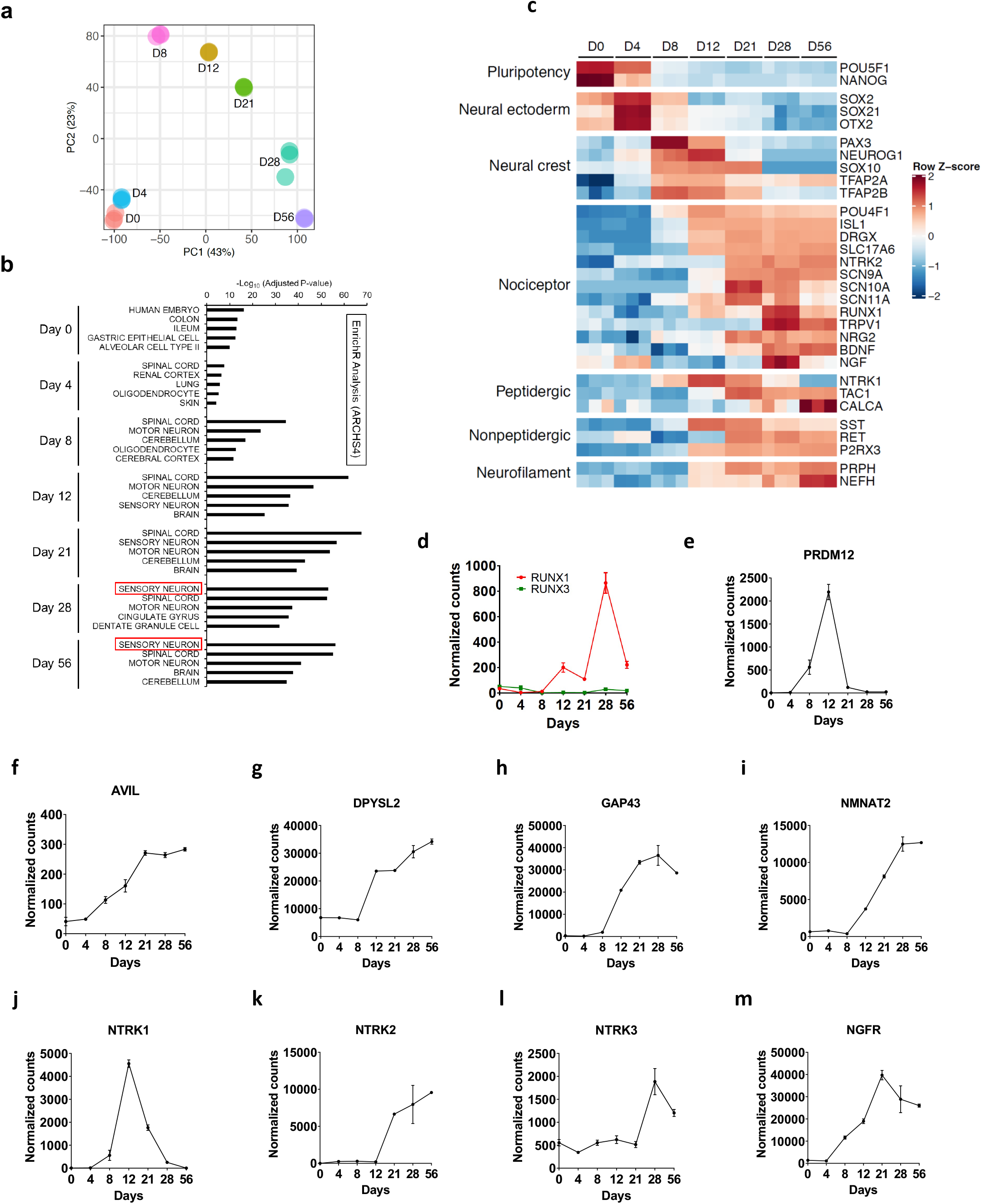
Transcriptomic analysis of the differentiation of hPSCs into neural crest and nociceptors. **a,** PCA plot of RNA-seq experiment showing distinct molecular signatures at each timepoint. **b,** Gene enrichment analysis using Enrichr and ARCHS4 database identify “sensory neuron” as top category at day 28 and 56. **c,** Heatmap analysis showing stepwise differentiation of pluripotent cells into neural crest and nociceptors (n = 3). **d-m,** Time-course expression profile of different genes with relevance in sensory neuron biology.

### Comparison of *in vitro*-generated neurons to human DRGs

To compare the gene expression signature of hPSC-derived nociceptors to *in vivo* tissues, we used datasets of human DRGs established by the GTEx consortium^34^. Pluripotent and differentiating cells, harvested at different timepoints, were compared to adult human DRG samples. Since pain is associated with structural and functional plasticity^7^, we also included tissue samples obtained from individuals with a clinical history of chronic pain. PCA plots showed that DRG samples clustered together and were distinct from cultured cells (**Fig. 3a**). However, while the pluripotent state was most distant from DRGs, differentiating cells followed a differentiation trajectory toward the DRG signatures. Hence, the transcriptome of nociceptors at day 28 and 56 were closest to DRG samples (**Fig. 3a**). Nociceptors are known to express a large numbers of ion channels and receptors^14^. To establish a resource for disease modeling and drug discovery, we performed comprehensive analyses comparing the expression of various gene families with special focus on ligand- and voltage-gated ion channels (potassium, sodium, calcium, chloride) and other cell membrane proteins (porins, gap junction proteins). This systematic comparison revealed a striking overlap of genes expressed by the induced nociceptors (day 28 and 56) and DRGs (**Fig. 3b-e and Extended Data Fig. 11**). The similarity of expressed genes in nociceptors and *in vivo* tissues provided additional evidence for the validity of this *in vitro* model.

**Fig. 3:**
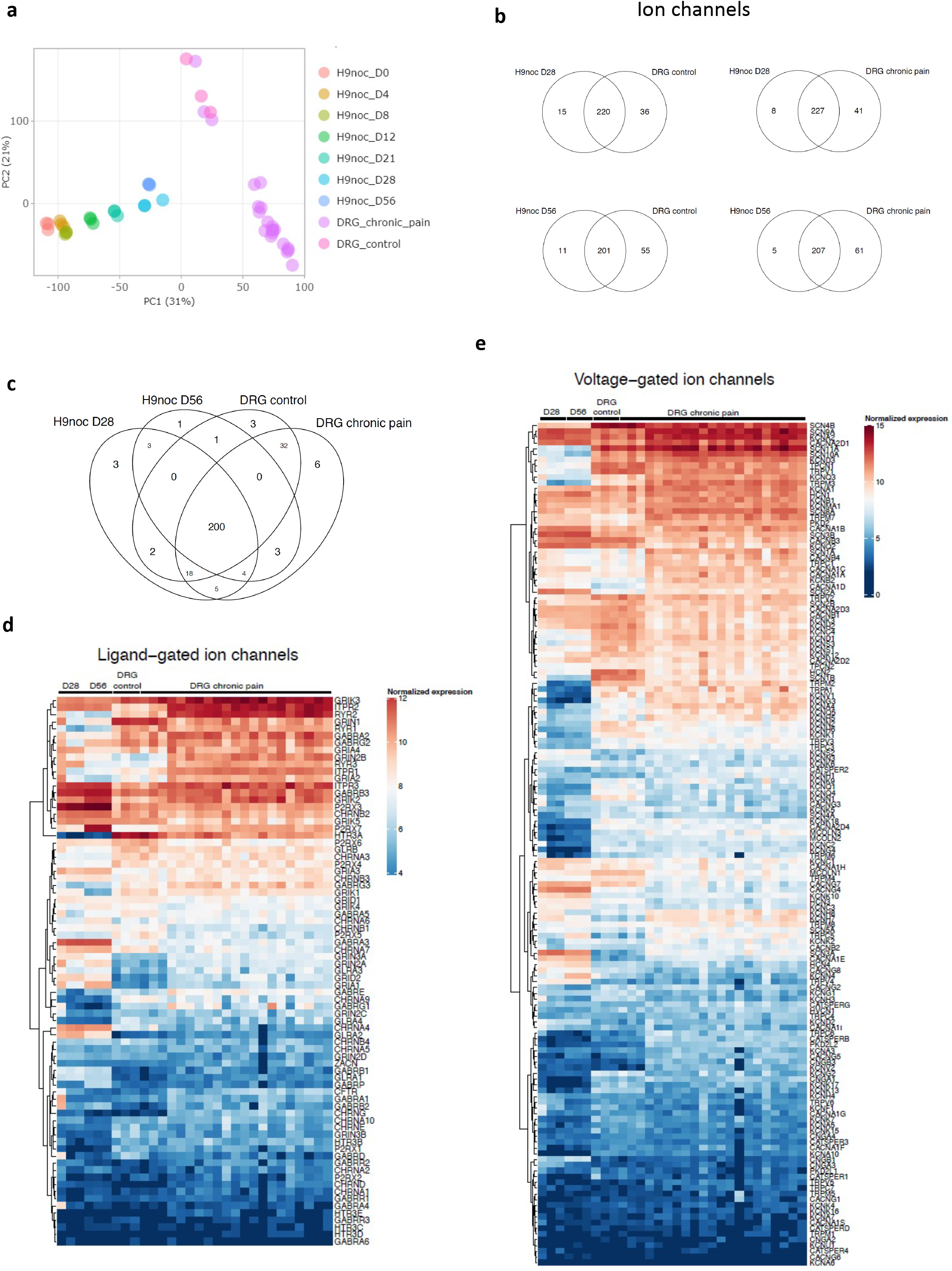
Molecular comparison of hPSC-derived nociceptors and human DRG samples. **a,** PCA of differentiating cells harvested at different timepoints and DRG samples (normal and chronic pain patient). **b, c,** Venn diagrams comparing expressed ion channels in nociceptors (day 28 and 56) and DRG samples. **d, e,** Heatmap comparison of ligand- and voltage-gated ion channels expressed by *in vitro*-generated nociceptors and DRGs (n = 3). See Extended Data Fig. 11 for detailed analysis of other gene families.

Next, we compared the transcriptome of nociceptors generated with our method with commercially available human iPSC-derived nociceptors (Axol Bioscience). Heatmap analysis showed similarity between day 28 nociceptors and commercial nociceptors (**Extended Data Fig. 12a**) but upon closer analysis, differences were found regarding the expression levels of specific genes including *RUNX1*, *NRG2*, *TAC1*, *PIEZO2*, *NGF*, *TRPC5* (**Extended Data Fig. 12b-h**). Although a large fraction of genes (n = 188) was found to be expressed both by sensory neurons generated with the present method as well as the commercially obtained cells (**Extended Data Fig. 13a**), 47 genes were exclusively expressed by the nociceptors generated with the present method. Conversely, 5 genes were expressed in commercial cells but were absent from our nociceptors. Comprehensive gene expression heatmaps comparing all detected channel genes were prepared for data mining (**Extended Data Fig. 13b-i**).

### Molecular targets for pain research and drug development

To generate hPSC-derived neurons for translational research, it is necessary to demonstrate consistent and reproducible derivation of large quantities of human nociceptors that express functional targets. We therefore performed a focused analysis of the transcriptome of hPSC-derived nociceptors and validated the expression of genes relevant for biomedical research. Using this approach, genes of interest expressed by nociceptors could be categorized into the following gene families: protein kinase, nuclear receptor, neurotransmitter transporter, ion channel, neuropeptide, GPCR (**Fig. 4a**). For instance, 502 protein kinases were expressed by nociceptors, which represents 73% of all protein kinases encoded by the human genome (**Fig. 4a**). Notably, the nociceptors expressed 44%, 48% and 22% of all known human neurotransmitter transporters, ion channels and neuropeptides, respectively (**Fig. 4a**). Moreover, the subcategories of expressed ion channels including sodium, chloride and calcium channels were measured as well (**Fig. 4b**). Next, we monitored the expression of other biologically diverse targets upregulated during the differentiation process using RNA-seq and immunocytochemical analysis Among these expressed targets were *P2RX3*, *TRPV1*, *TAC1*, *CALCA*, Na_V_1.7 (*SCN9A*), Na_V_1.8 (*SCN10A*) and the nociceptin receptor *OPRL1* (**Fig. 4c-h and Extended Data Fig. 14**). Expression of the sodium channel Na_V_1.9 (SCN11A) was detected on the transcript level (**Extended Data Fig. 14a**) and additional experiments using quantitative *in situ* hybridization (RNAscope) provided insights into Na_V_1.7, 1.8 and 1.9 expression and co-localization at the cellular level (**Fig. 4j,k**)

**Fig. 4:**
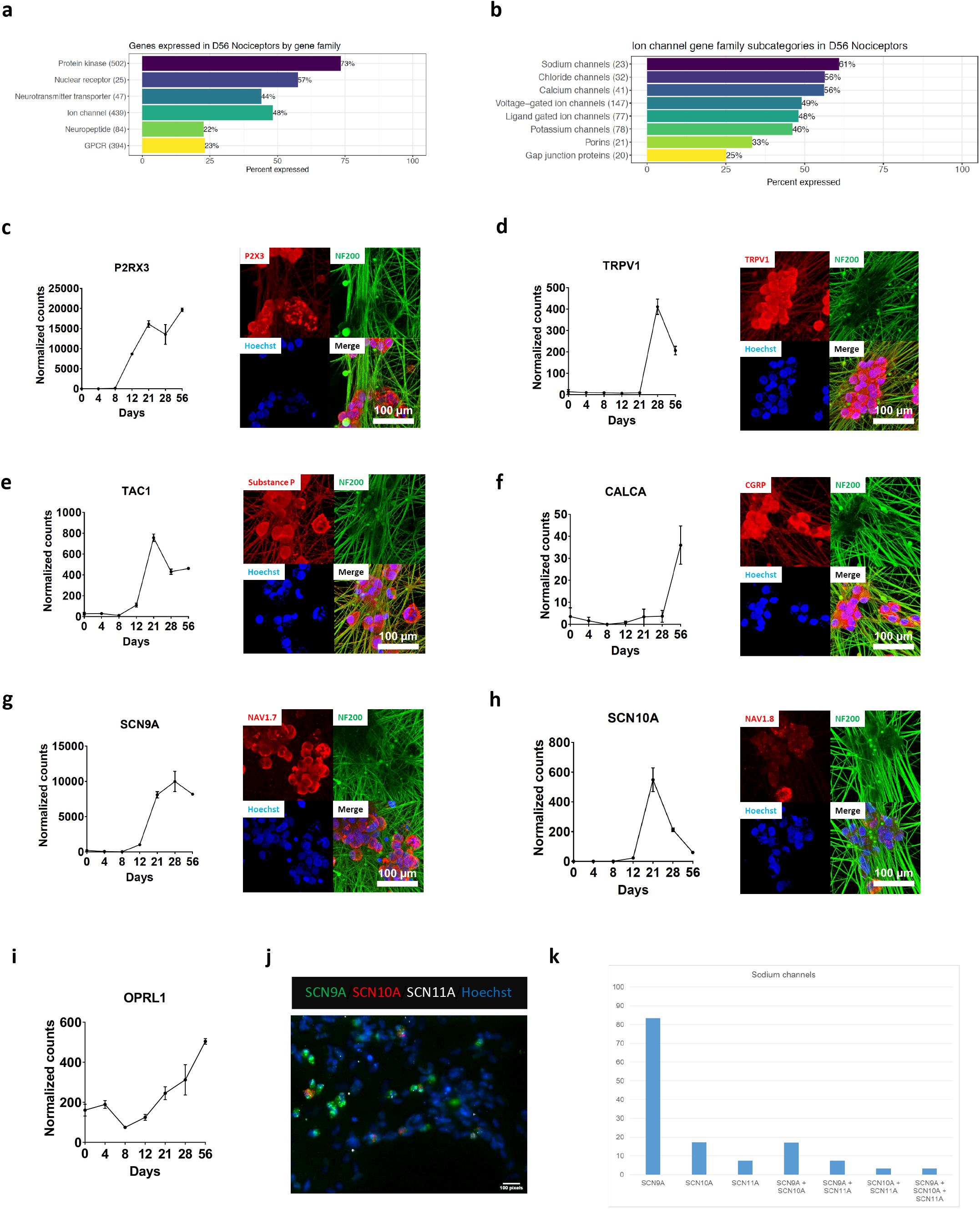
Analysis of gene families and specific targets expressed by hPSC-derived nociceptors a, b, Overview of genes and gene families expressed by hPSC-derived nociceptors (RNA-seq). **c-i**, Expression of major targets by nociceptors, as confirmed at the transcript and protein level (RNA-seq and immunocytochemistry)(n = 3, mean ± s.d.). **j, k,** Quantitative *in situ* hybridization (RNAscope) and analysis of sodium channel expression and co-expression of Na_v_ 1.7, 1.8 and 1.9 in nociceptors.

### Functional characterization of nociceptors

To investigate the functional properties of hPSC-derived nociceptors, we performed experiments using various methods, specific agonists, and stimulation protocols. First, calcium imaging was carried out using a cellular screening platform and nociceptors were exposed to DMSO (control), 40 mM KCL, 10 µM ATP, 1 µM capsaicin, 100 µM mustard oil, and 250 µM menthol (**Fig. 5a**). Reagent concentrations used in these experiments were chosen to ensure that only the respective cognate receptors were activated^21,35^. Neuronal depolarization with KCL, which activates voltage-gated calcium channels, elicited the strongest signal (**Fig. 5a**). Differential cellular response was observed after activation of specific receptors such as P2RX3 (ATP), TRPV1 (capsaicin), TRPA1 (mustard oil), and TRPM8 (menthol)(**Fig. 5a**). Additional functional experiments using an automated multi-electrode array system confirmed that nociceptors were responsive to α,β-methylene-ATP, capsaicin, and mustard oil (**Fig. 5b**). In an independent experiment, when the temperature was increased in the MEA system from 37 °C to 40 °C, enhanced neuronal excitability was detected, suggesting the presence of temperature-sensitive receptors (**Fig. 5c,d**). A typical feature of nociceptors is sensitization, which is a mechanism that contributes to the development of clinical pain^36^. To test whether hPSC-derived nociceptors could be sensitized similar to a previous report^37^, we treated cells with the chemotherapeutic drug oxaliplatin and the inflammatory mediator PGE2 (prostaglandin E2) for 10 min (**Fig. 5c, d**). To stimulate nociceptors, the temperature was then increased in the MEA system from 37 °C to 40 °C. Cells treated with oxaliplatin and PGE2 were more excitable than DMSO controls, indicating that nociceptors could be sensitized in this *in vitro* assay (**Fig. 5c, d**).

**Fig. 5:**
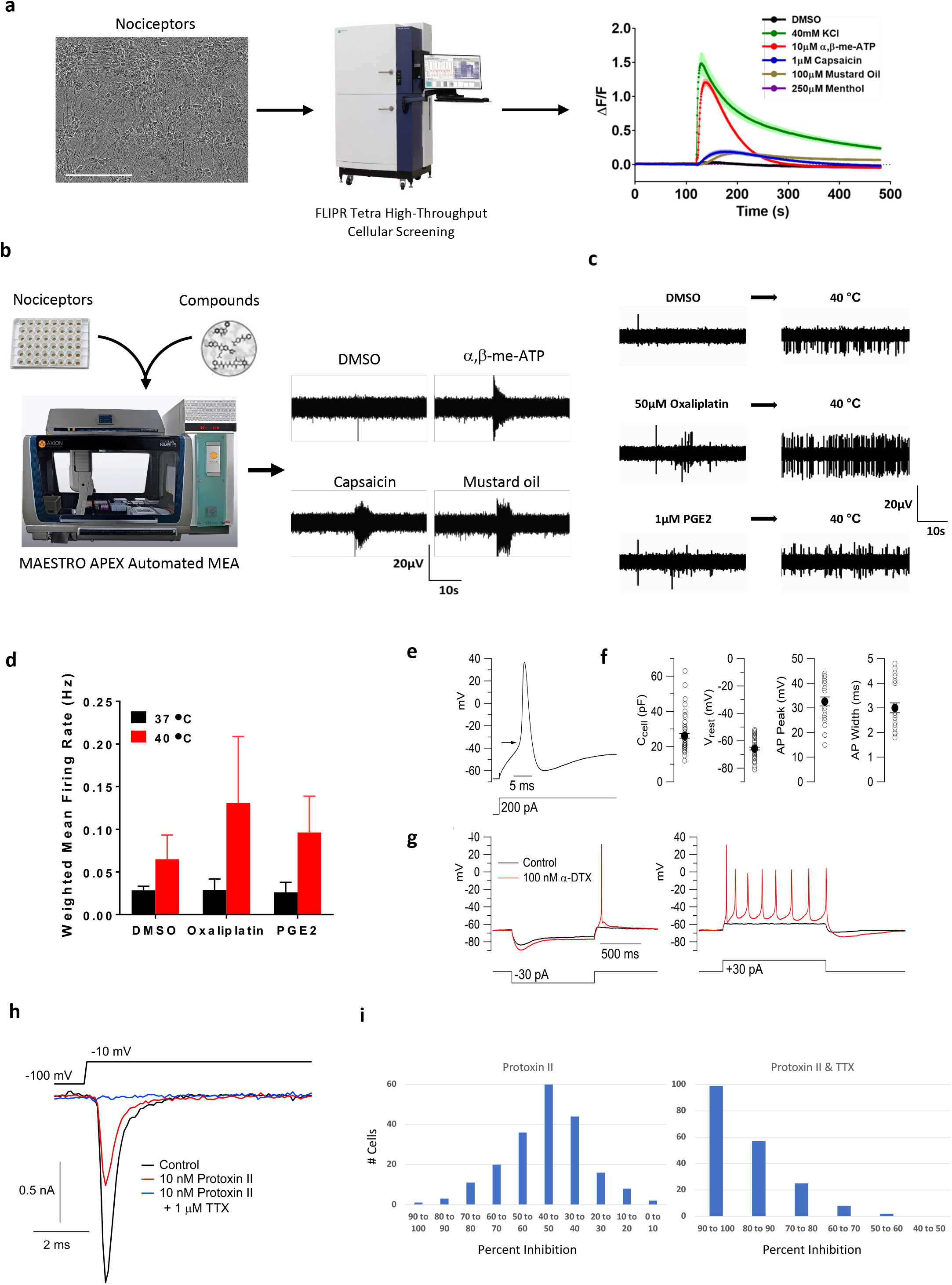
Functional characterization of hPSC-derived nociceptors. **a,** Calcium flux analysis after stimulation of nociceptors with KCL, ATP, capsaicin, mustard oil and menthol (n = 8, mean ± s.d.). **b,** Automated MEA showing that nociceptors are activated by specific ligands. **c, d,** MEA experiment showing that nociceptors are stimulated by a temperature increase from 37 to 40°C and are presensitized after treatment with oxaliplatin or PGE2 (n = 3, mean ± s.d.). **e,** Typical action potential in nociceptors evoked by current injection. Arrow indicated the threshold voltage (- 35 mV). **f,** Collected results for passive and active electrical properties of the neurons. Open circles show measurements from individual cells and close circles show mean ± s.e.m. (n = 53 cell capacitance and resting potential; n = 25 for action potential peak and width). **g,** Enhancement of excitability by the K_v_1-inhibitor *α*-dendrotoxin (DTX). **h,** Automated single-cell patch clamp (Sophion Qube) record showing effect of 10 nM Protoxin-II and 1 μM TTX (in the continuing presence of Protoxin-II) on sodium current in an hPSC-derived nociceptor. **i,** Collected results for percentage inhibition of sodium current by Protoxin-II alone and by Protoxin-II plus TTX, in 217 neurons.

To further investigate the electrical properties of hPSC-derived sensory neurons, we performed recordings using patch clamp techniques (**Fig. 5e-g**). Neurons had well-polarized resting potential, with an average value of −65.7 ± 1.0 mV (mean ± s.e.m., n = 53) and had relatively high input resistances (391 ± 23 MOhms, n = 53), indicative of cell health. Almost all neurons (25 of 26) tested in current clamp mode with current injections of increasing size fired robust action potentials, with an average action potential height of 97 ± 2 mV (n = 25), reaching a peak voltage of + 33 ± 2 mV (n = 25), with an average width of 3.0 ± 0.2 ms (measured at half-amplitude, n = 25). Most neurons (20 of 25) showed strong adaptation of action potential firing, firing a single action potential in response to current injections of up to 300 pA, similar to a subset of mature mouse primary nociceptors^38^. As with many native mouse DRG neurons, the strong adaptation appears to be conferred in part by expression of Kv1 family potassium channels, which are highly effective in limiting firing in response to maintained current injections to a single action potential^38^. As with rodent nociceptors displaying this characteristic, the Kv1 inhibitor *α*-dendrotoxin enhanced the excitability of the neurons, decreasing the threshold current needed to evoke an action potential and causing repetitive firing during current injections (**Fig. 5g**). Enhanced repetitive firing and decreased rheobase current were seen in 4 of 5 cells tested with 100 nM *α*-dendrotoxin.

Nociceptor-selective sodium channels are promising targets to develop new analgesics^7,39^. To test whether hPSC-derived nociceptors could be used in high-throughput fashion, we performed a functional screen using an automated patch-clamp system (Qube 384, Sophion Bioscience). To this end, day-28 neurons were dissociated and plated into single-hole 384-well QChips for patch-clamp recording. Voltage-dependent sodium currents were evoked by voltage steps to −10 mV from a holding potential of −100 mV. Application of 10 nM Protoxin-II (ProTx-II), a selective inhibitor of voltage-gated Nav1.7 channels^40^, inhibited an average of 46.2% ± 15.7% (mean ± s.d.; n = 201) of the total sodium current and with subsequent application of 1 μM TTX together with Protoxin-II, an average of 88.5% ± 9.8% (mean ± s.d., n = 191) of the total initial sodium current was inhibited (**Fig. 5h,i**).

### Elucidating drug effects using human nociceptors

To further demonstrate translational utility of hPSC-derived nociceptors, we asked whether specific drugs could be studied in relevant assays. Such experiments are important because they provide information about the quality of human cells and molecular targets of interest, but also help characterize the efficacy and safety of established drugs or drug candidates in preclinical development. First, we tested several commercially available P2RX3 antagonists, to find out whether it might be possible to identify the most potent inhibitor using a phenotypic readout. P2RX3 is a subtype of ionotropic purinergic receptors expressed specifically by nociceptors and involved in nociceptive transmission^41^. Strong expression of P2RX3 was detected by RNA-seq, as described above (**Fig. 4c**). The human nociceptors were pretreated with DMSO (control), four selective and potent inhibitors of P2RX3 (RO-51, TNP-ATP, TC-P 262, RO-3) and a non-selective P2 purinergic antagonist (PPADS) for 30 minutes, followed by stimulation with α,β-methylene-ATP, a specific purinergic receptor agonist. Automated MEA analysis revealed that RO-51 was the most potent inhibitor completely blocking the effect of α,β-methylene-ATP in hPSC-derived nociceptors (**Fig. 6a**).

**Fig. 6:**
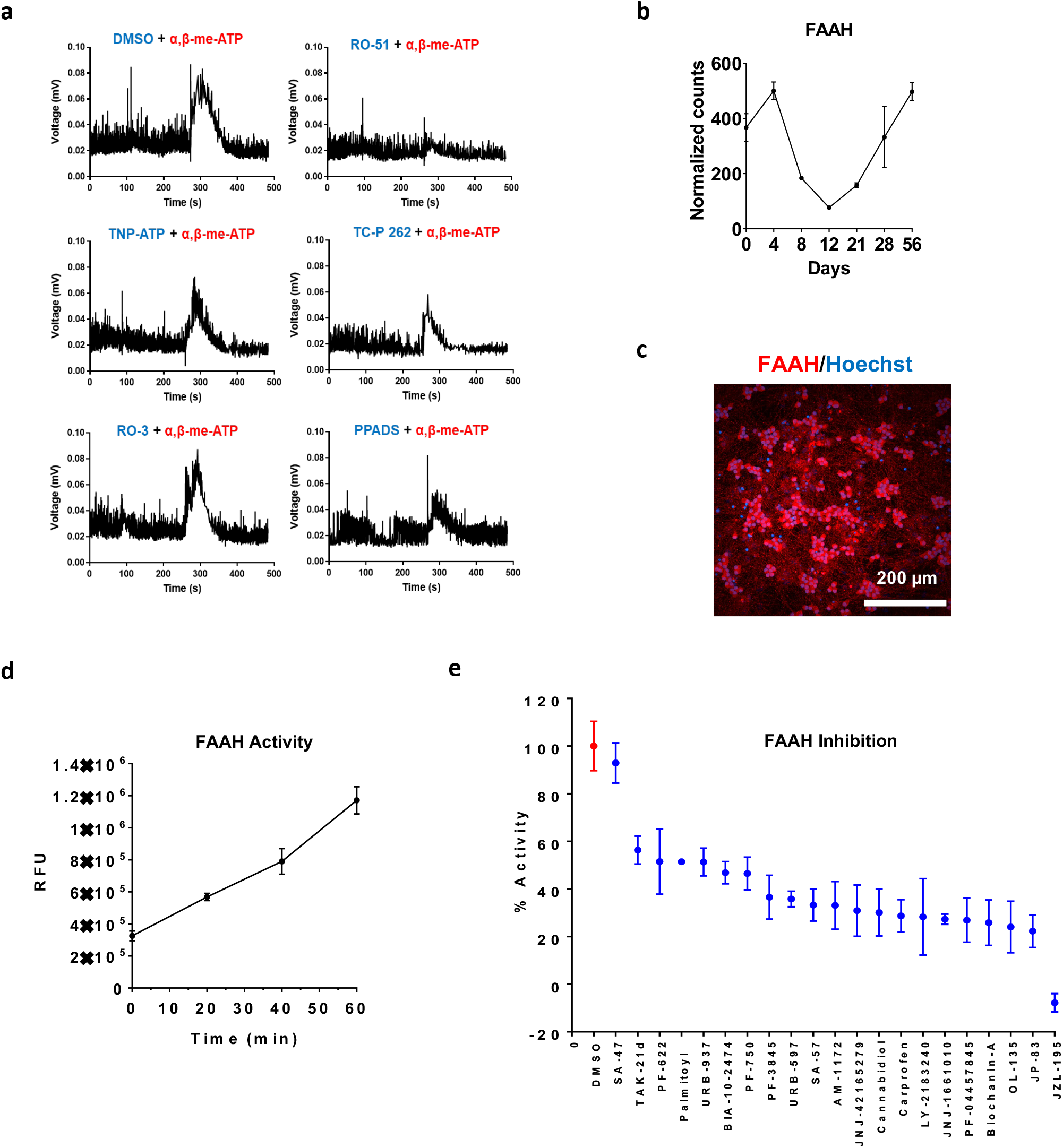
Drug testing using hPSC-derived nociceptors. **a,** MEA experiment comparing the potency of various P2RX3 inhibitors in human nociceptor cultures. **b,** Gene expression of FAAH during cell differentiation into nociceptors (RNA-seq) (n = 3, mean ± s.d.). **c,** Immunocytochemical analysis of FAAH expression in nociceptors (day 28). **d,** Real-time fluorescent assay monitoring FAAH activity in nociceptors (n = 8, mean ± s.d.). **e,** FAAH inhibition assay testing the relative efficacy of 21 known inhibitors in day-28 nociceptor cultures (n = 8, mean ± s.d.).

Rare genetic mutations that cause congenital insensitivity to pain can inform about new drug targets^2^. Fatty acid amide hydrolase (FAAH) is a critical enzyme responsible for breakdown of several bioactive lipids such as anandamide, which act as endogenous ligands for cannabinoid receptors. FAAH knockout mice and patients with hypomorphic SNPs (single nucleotide polymorphisms) or mutations show hypoalgesia^42^. Therefore, inhibition of FAAH emerged as a target for modulating pain but clinical trials have been unsuccessful thus far^43,44^. Using the hPSC-derived nociceptors, we first confirmed that FAAH was expressed at the transcript and protein level (**Fig. 6b,c**). Next, we established an enzymatic assay to evaluate how FAAH catalyzes the hydrolysis of AAMCA (Arachidonyl 7-amino, 4-methly coumarin amide) into arachidonic acid and AMC (7-Amino 4 methly coumarin)^45^. Because AMC is fluorescent, the activity of FAAH can be continuously monitored over time in the human neurons (**Fig. 6e**). Using this screening-compatible assay, we tested a panel of known FAAH inhibitors, including those that were unsuccessful in clinical trials (e.g., PF-04457845, BIA 10-2474, JNJ-42165279). When these compounds were administered to day 28 nociceptors, JZL-195 was identified as most efficient in blocking FAAH activity. Interestingly, SA-47 reported to be a selective and potent FAAH inhibitor in rodent cells^46^ showed poor efficacy in this assay. All the other inhibitors showed differential effects on FAAH activity (**Fig. 6f**). This screen demonstrated that the *in vitro*-generated human nociceptors can play a role in drug development since they express cell type-specific targets and enable the characterization of drugs, which offers new opportunities for human biology-based phenotypic drug screens.

## Discussion

Development of new analgesics has largely failed over the last decades and the current opioid crisis in the United States underscores the need for effective and safe pain-relieving drugs without abuse liability, tolerance, and other adverse effects^7^. Generation of well-characterized human nociceptors from iPSCs as an inexhaustible source is urgently needed for basic and translational pain research and drug development. Here, we developed a new method for the efficient production of sensory neurons from hPSCs. These *in vitro*-generated neurons were extensively characterized and displayed the unique structural, molecular, and functional properties of native sensory neurons of the DRG. Importantly, nociceptors showed robust expression of various molecular targets and ion channels and screening experiments demonstrated utility in translational assays relevant for drug testing. The extensive datasets and systematic comparisons to DRG neurons establish a unique resource and should also leverage disease modeling and toxicology studies (e.g., genetic disease, chemotherapy-induced neuropathy).

Furthermore, it is remarkable that neurons with pseudo-unipolar morphology can be generated *in vitro* since positional molecular cues and secreted factors from the microenvironment, including chemo-attractants and chemorepellents, that typically guide axonal growth and pathfinding in the developing embryo are absent *ex vivo*. Based on the presented data, it appears conclusive that induction of the appropriate genetic program leads to a cell-intrinsically determined polarization of induced neurons resembling DRG neurons. A previous report ectopically expressed Brn3a with either Ngn1 or Ngn2 in human and mouse fibroblasts and reported the emergence of pseudo-unipolar sensory neurons^20^. However, reprogramming efficiency was low and definitive evidence for axon splitting and ultrastructural analysis was not provided. Similarly, other studies describing differentiation of hPSCs into sensory neurons^13,15,19^ did not report the generation of neurons with distinct pseudo-unipolar anatomy. The structure-function relationship is particularly important for polarized neuronal cells and expression and localization of ion channels have distinctive patterns in cell bodies and different axon segments (e.g., proximal versus distal, peripheral versus central projection). Currently, little information is available on ion channel distribution in the DRG system^7^. Streamlined production of pseudo-unipolar sensory neurons from human iPSCs should help to better understand ion channel biology and neuronal excitability with the goal to develop nociceptor-selective analgesics. Lastly, the robustness and scalability of the human sensory neuron model presented here should also pave the way for elucidating the precise molecular mechanisms of sensory neuronal axonal degeneration and regeneration in future studies.

## Supporting information

Ext Data Fig 1

Ext Data Fig 2

Ext Data Fig 3

Ext Data Fig 4

Ext Data Fig 5

Ext Data Fig 6

Ext Data Fig 7

Ext Data Fig 8

Ext Data Fig 9

Ext Data Fig 10

Ext Data Fig 11

Ext Data Fig 12

Ext Data Fig 13

Ext Data Fig 14

## Acknowledgements

We thank Paul Shinn, Misha Itkin, Zina Itkin, Carleen Klumpp-Thomas, John Braisted, Charles Bonney, Yeliz Gedik, and Steve Pittenger for their support throughout this work. We are grateful to Alan Hoofring and Ethan Tyler from the NIH Medical Arts Design Section for their technical expertise. We acknowledge the excellent work by Weifeng Yu, Mei Zhang, and Sung Hoon Park at Sophion Bioscience.

## EXTENDED DATA FIGURE LEGENDS

**Extended Data Fig. 1: Hotspot kinase assay identifies CHIR98014 as superior WNT pathway activator**

**a,** Direct comparison of two GSK3beta inhibitors reveals that CHIR98015 has greater potency than the commonly used CHIR99021 (n = 2, representative data points presented). **b,** Comprehensive profiling was performed to individually inhibit a panel of 369 human wild-type kinases using 1.5 µM CHIR99021 and 0.5 µM CHIR98014. Note that target specificity is similar for both compounds but CHIR98014 is more potent. Different colors in the tree depict different kinase families. For more details see also Supplementary Table 1.

**Extended Data Fig. 2: Sphere formation promotes neural crest specification**

**a,** Monolayer differentiation with CHIR98014 and A83-01 promotes spontaneous sphere formation (arrows) and expression of the neural crest marker SOX10 and the sensory neuron marker ISL1. Note that adherent cells surrounding the spheres do not express SOX10. **b,** Manual transfer and attachment of spheres shown in (a) on Geltrex results in cell migration and neuronal differentiation. **c,** Observations mentioned above (a, b) led to protocol development for neural crest and nociceptors by introducing a differentiation stage based on sphere formation (for more details see also Fig. 1a).

**Extended Data Fig. 3: Establishing PD173074 as superior inhibitor of FGF signaling**

**a,** Inhibition of FGFR1 by PD173074 and SU5402 (n = 2, individual data points presented). **b,** Hotspot kinase profiling was used to individually inhibit a panel of 369 human wild-type kinases using 0.5 µM SU5402 and 25 nM PD173074. Note that SU5402 inhibits TrkB and TrkC receptors with even higher potency than FGF receptors. In contrast, PD173074 specifically inhibits FGFR1/2/3 and has no off-targets. Different colors in the tree depict different kinase families. For more details see also Supplementary Table 1.

**Extended Data Fig. 4: Monitoring expression of SOX10 during cell differentiation a,** Gene expression profile of *SOX10* (RNA-seq) is consistent with neural crest differentiation trajectory and nociceptor generation (n = 3, mean ± s.d.).

**b,** Immunocytochemical analysis of the proliferation marker Ki-67, the neural crest marker SOX10, and the nociceptor marker BRN3A. Note that the number of nociceptors increase from day 15-28 at the expense of cells expressing Ki-67 and SOX10, consistent with increased cell differentiation and maturation over time.

**Extended Data Fig. 5: Robotic cell culture and production of nociceptors**

**a,** Overview image of the CompacT SelecT platform. **b,** iPSCs and differentiating cells, including nocispheres, are cultured in large bar-coded T175 flasks by the CompacT SelecT system. A typical workflow generates over 150 million nociceptors (30 cryopreservable vials), which can be scaled up 30-fold depending on experimental needs such as parallel-processing of multiple cell lines.

**Extended Data Fig. 6: Immunocytochemical analysis of nociceptors**

**a,** Expression of the axonal protein TAU (MAPT) by BRN3A+ nociceptors. **B, c,** Nociceptors are immunoreactive against vGLUT1 and glutamate indicating their glutamatergic neurotransmitter phenotype. **D,** Neuronal cells with immunopositivity for the inhibitory neurotransmitter GABA were only found very sporadically. **e,** Cells expressing tyrosine hydroxylase (TH) were not detected. **f,** Pseudo-unipolar nociceptors develop prominent NF200^+^ axons, and MAP2 stains cell bodies only, due to the lack of any dendritic structures.

**Extended Data Fig. 7: Confocal microscopic analysis showing axon splitting**

**a, b,** Immunostaining for neuronal marker TUJ1 and Ankyrin G, a protein enriched at the axon initial segment. Arrow shows a neuron displaying axon splitting. **c,** Confocal sections (z-stack images) of the same neuron shown in a and b.

**Extended Data Fig. 8: Ultrastructural analysis of peptidergic and non-peptidergic neurons**

**a, b**, Examples of peptidergic neurons with prominent cytoplasmic LDCVs (red arrows). Overview of the cell body (b) is presented in Fig. 1i. **c,** Axonal structures containing multiple LDCVs (red arrows). **d,** Example showing non-peptidergic nociceptors with round nuclei and well-developed cytoplasmic organelles. **e, f,** Axonal profiles sectioned in different planes containing mitochondria but which lack LDCVs. Scale bar, 2 µm.

**Extended Data Fig. 9: Ultrastructure of axon initial segment**

**a,** Overview image showing nociceptor cell bodies and axonal structures and LDCVs. White arrows indicate the axon initial segment, which emanates from the cell soma. Red arrows point to more distal portions of the same axon. **b,** Higher magnification of the axon shown in (a). Sale bar, 2 µm.

**Extended Data Fig. 10: Expression profile of genes reflecting differentiation of pluripotent cells into sensory neurons**

**a-g,** Time-course of genes, expressed and downregulated at specific timepoints, indicate stepwise differentiation of pluripotent cells (POU5F1/OCT4; NANOG) into glutamatergic nociceptors. **h,** Genome browser showing expression of specific gene transcripts, consistent with generation of nociceptors from pluripotent cells.

**Extended Data Fig. 10: RNA-seq analysis of genes reflecting differentiation of pluripotent cells into neural crest and sensory neurons**

**a-f,** Time-course of genes, expressed and downregulated at specific timepoints, indicate stepwise differentiation of pluripotent cells (*POU5F1*/*OCT4*; *NANOG*) into glutamatergic nociceptors (n = 3, mean ± s.d.). **g,** Genome browser showing expression of specific gene transcripts, consistent with generation of nociceptors from pluripotent cells (italicize gene names). **h-i,** Strong induction and distinct gene expression profiles of *GFRA1*, *GFRA2*, and *GFRA3* (n = 3, mean ± s.d.).

**Extended Data Fig. 11: Heatmap analysis of hPSC-derived nociceptors and DRG transcriptomes**

Comparison of expressed transcripts for various ion channels (potassium, sodium, chloride, calcium), porins and gap junction proteins. Note the high degree of similarity between *in vitro*-generated nociceptors and DRGs from healthy individuals and patients with chronic pain.

**Extended Data Fig. 12: RNA-seq and comparison**

**a,** Heatmap analysis of genes expressed by nociceptors (day 28) generated with the current method versus Axol Bio neurons (n = 3).

**b-h,** Direct comparison of transcripts of single genes reveals different expression levels for *RUNX1*, *NRG2*, *TAC1*, *PIEZO2*, *NGF*, *TRPC5*, and *TRPV1* (n = 3, mean ± s.d.).

**Extended Data Fig. 13: Ion channel expression in hPSC-derived nociceptors and commercially obtained nociceptors (Axol Bio)**

**a,** Venn diagram showing the expressed ion channel genes in nociceptors generated with the present method and commercially obtained neurons (Axol Bio). **b-i,** Comparative heatmap analysis of various gene families and ion channels as indicated.

**Extended Data Fig. 14: Expression of genes with relevance for translational pain research**

**a, b,** Time-course RNA-seq experiment and raw data of expressed transcripts in three different replicates per timepoint showing consistency of directed differentiation.

## METHODS

### hPSC Cell culture

All hESCs (WA09; WiCell) and hiPSCs (LiPSC-GR1.1, NCRM5, CMT2A1.1, CMT2A2.1. CMT2A3.1) were maintained under feeder-free conditions in Essential 8 (E8) cell culture medium (Thermo Fisher Scientific) and vitronectin (Thermo Fisher Scientific) coated microplates or T175 flasks. Cells were passaged every three days. The hPSC colonies were treated with 0.5 mM EDTA in phosphate buffered saline (PBS) without calcium or magnesium (Thermo Fisher Scientific) for 5-6 min to dissociate the hPSC colonies. The resulting cell clumps were counted using the Nexcelom Cellometer automated cell counter. The clumps were then plated at a density of 1.5 × 10^5^ cells per cm^2^ in E8 cell culture medium supplemented with the CEPT cocktail^22^ for the first 24 h. The CEPT cocktail, consisting of 50 nM Chroman 1 (#HY-15392; MedChem Express), 5 μM Emricasan (#S7775; Selleckchem), Polyamine supplement (#P8483, 1:1000 dilution; Sigma-Aldrich), and 0.7 μM Trans-ISRIB (#5284; Tocris), was used to improve cell survival and provide cytoprotection during cell passaging. Cells were maintained in a humidified 5% CO_2_ atmosphere at 37°C.

### Differentiation of hPSCs into nociceptors

hPSC maintained in E8 medium on VTN-coated 6-well plates, as described above, were dissociated using 0.5 mM EDTA for 5-6 min. The resulting clumps were then counted using the Nexcelom Cellometer and seeded at a density of 1.5 x 10^5^ cells/cm^2^ in E8 medium on VTN-coated 6-well plates. 24 h later (Day 0), the spent medium was replaced with daily with E6 medium containing 2 μM A83-01 and 0.2 μM CHIR98014 (designated as Noci-1 medium). On Day 3, Noci-1 medium was aspirated, cells were washed with PBS and incubated in 1 mL Accutase per well (6-well plates) for 5 min at 37°C. Cell were resuspended in 5 mL Noci-2 medium (see Fig. 1a) and dissociated into single cells by gently pipetting up and down 10 times, followed by centrifugation at 300 g for 3 min at room temperature. Supernatants were removed and cells were resuspended in Noci-2 medium + CEPT and plated at 5.5 x 10^6^ cells/well into 6-well AggreWell 800 plates according to manufacturer’s instruction (STEMCELL Technologies) for free-floating suspension culture and nocisphere formation. For Day 4-14 (nocisphere stage), perform daily medium change with Noci-2 medium (no CEPT required). Spent medium (4mL) was gently removed and fresh medium was added against the wall to avoid disturbing the nocispheres. At Day 14, one 6-well plate with nocispheres was collected into 50 mL tubes, allowed to settle down for 5 minutes, cell culture medium was removed and spheres were wash with 10 mL PBS and allowed to settle down again. After removal of PBS, spheres were dissociated using the MACS EB Dissociation Kit (order no. 130-096-348) according to manufacturer’s instruction (Miltenyi Biotec). The cell pellet was resuspended using Noci-3 medium (see Fig. 1a) + CEPT and pass the cells through 70 µm cell strainer. Cells were counted and plated at 2 x 10^6^ cells/well into Geltrex coated 6-well plates using Noci-3 medium + CEPT (remove CEPT after 24 hours). The cells can also be cryopreserved at this time point at 5 million cells/mL using Noci-3 medium + CEPT + 10% DMSO. Noci-3 medium was changed every 2-3 days until Day 28 or longer if further maturation is needed.

### Automated cell culture

Scalable robotic cell culture and differentiation was carried out by using the CompacT SelecT platform as previously described^25^ and following the steps described above. For sphere formation, T175 flask were pretreated with anti-adherence solution (STEMCELL Technologies).

### Immunocytochemistry

hPSC and hPSC-derived nociceptors at different time points were cultured as described above on glass-bottom multiwell plates (P24-1.5H-N, Cellvis). The cultures were fixed with 4% formaldehyde (28908, Thermo-Fisher) in PBS for 20 min, followed by permeabilization with 0.2% Triton X-100 (85111, Thermo-Fisher) in PBS for 10 min. Then the cultures were incubated with PBS + 1% bovine serum albumin (A9418, Millipore-Sigma) for 1 h at room temperature, followed by incubation with primary antibodies overnight at 4°C. Secondary antibodies were incubated at room temperature for 1 h. Then the cultures were stained with 4 µM Hoechst 33342 (PI62249, VWR) in PBS for 10 min before imaging on Opera Phenix high-content microscope (PerkinElmer) or Zeiss LSM 710 confocal microscope. Primary and secondary antibodies used are summarized in Supplementary Table 2.

### Electron Microscopy

Day 28 nociceptors cultured on Geltrex-coated 6-well plate are processed, embedded, and made thin-sections using in situ methods for the EM analysis. Briefly, cells are fixed in 2% glutaraldehyde (v/v) in sodium cacodylate buffer (0.1M, pH 7.4) followed by rinsed in cacodylate buffer, and post-fixed by 1% osmium tetroxide (v/v) (Electron Microscopy Sciences) in cacodylate buffer. Cells are rinsed in cacodylate buffer, then in acetate buffer (0.1M, pH 4.0) prior to uranyl acetate (0.5% w/v, pH 4.5) en bloc stained. Cells are dehydrated in series of ethanol (e.g., 35%, 50%, 75%, 95%, and 100%) are washed three changes in pure epoxy resin (Electron Microscopy Sciences) and infiltrated overnight. Cells are embedded in a fresh epoxy resin following day and cured in 55 °C oven for 48 hours. The cured epoxy in the 6-well plate is separated by submerged in liquid nitrogen. Thin sections (60 to 70nm) are made in parallel direction to the growth of cells using a diamond knife and ultramicrotome (Leica). Thin sections are mounted on 150 copper mesh grids and counter stained in aqueous uranyl acetate (0.5% w/v) and Reynold’s lead citrate. The sections are examined using an electron microscope (Hitachi) and images were capture by using a CCD camera (AMT).

### Bulk RNA-seq

hPSC and hPSC-derived nociceptors at different time points were lysed using buffer RLT+ (1053393, Qiagen) supplemented with 2-mercaptoethanol (63689, Millipore-Sigma) directly in wells and RNA was extracted and purified using RNeasy Plus Mini Kit (74136, Qiagen) according to the manufacturer’s instruction. QIAcube automated workstation was used for the extraction (Qiagen). Genomic DNA was eliminated by both the gDNA eliminator column and on-column incubation with DNase I (79256, Qiagen). Sequencing libraries were constructed and sequenced at the National Cancer Institute’s Center for Cancer Research sequencing core facility using Illumina TruSeq® Stranded mRNA kit.

### Multi-electrode array (MEA)

Nociceptor activity was analyzed using the APEX robotic maestro multi-electrode array system (Axion Biosystems) according to the manufacturer’s protocol. In Brief, Day 14 nociceptors were thawed in Noci-3 medium + CEPT and plated at a density of 80,000 cells/well on PLO/Laminin coated 48-well MEA plates. Twenty-four hours post-plating the plating medium was replaced by Noci-3 medium + 1 μM PD0332991 and full medium change was performed every 2-3 days. Nociceptors were recorded 14 days post plating and 10 μM α,β-me-ATP, 5 μM capsaicin or 100 μM mustard oil were used to stimulate the cells. For sensitization assay, nociceptors were pre-treated with 50 μM Oxaliplatin or 1 μM PGE2 for 30 minutes and then recording temperature was increased from 37°C to 40°C to record the thermo responses. For P2X3 inhibitory assay, nociceptors were pre-treated with 10 μM of RO-51, TNP-ATP, TC-P 262, RO-3 or PPADS for 30 minutes and stimulated with 10 μM α,β-me-ATP.

### Patch clamp

Recordings were performed at room temperature using a Multiclamp 700B Amplifier (Molecular Devices). Voltage or current commands were applied and signals were recorded using a Digidata 1321A data acquisition system (Molecular Devices) controlled by pCLAMP 10.3.2 software (Molecular Devices). Electrodes were pulled on a Sutter P-97 puller (Sutter Instruments) and shanks were wrapped with Parafilm (American National Can Company) to allow optimal series resistance compensation without oscillation. The resistances of the pipettes were 2-3.5 MΩ when filled with the internal solution consisting of 140 mM K aspartate, 13.5 mM NaCl, 1.6 mM MgCl_2_, 0.09 mM EGTA, 9 mM HEPES, 14 mM creatine phosphate (Tris salt), 4 mM MgATP, 0.3 mM Tris-GTP, pH 7.2 adjusted with KOH. Seals and recordings were made in an external Tyrode’s solution consisting of 155 mM NaCl, 3.5 mM KCl, 1.5 mM CaCl_2_, 1 mM MgCl_2_, 10 mM HEPES, 10 mM glucose, pH 7.4 adjusted with NaOH. After establishing whole-cell recording, cell capacitance was nulled and series resistance was partially (70-85%) compensated. Reported membrane potentials are corrected by −10 mV from apparent voltages to account for the junction potential of the internal solution relative to the external solution when zeroing the electrode current before recording.

Cell capacitance was measured from 5 mV hyperpolarization from −80 to −85 mV, subtracting currents immediately before and after breaking through the membrane into whole-cell mode in order to separate cell capacitance from any pipette capacitance that was not fully compensated before break-through. Sequence of inhibitors for dissecting K current components. Resting potential and input resistance were measured either from a series of stairstep voltage changes from −85 to −35 mV delivered while breaking-through into whole cell mode (with resting potential measured as the voltage at which current became zero) or after switching the amplifier into current clamp mode, using hyperpolarizing or small depolarizing current steps to define input resistance. Threshold voltage was defined as the voltage at which the derivative of voltage reached 4% of the maximum upstroke velocity of the action potential.

### Automated patch clamp

Sensory neurons (day 28) were dissociated as follows: cells were first washed 2 times with PBS, and then were treated with 3∼4 mL 0.05% Trypsin per T175 cell flask for approximately 7 min at 37°C. Subsequently, 3∼4 mL culture medium with FBS was added and then cells were triturated 10 times with 5 ml pipette. After that, cells were then spun down (2 min with 1000 rpm) and resuspended in 6 ml serum-free medium (Sigma Cat#: C5467 with 25 mM added HEPES, P/S, and 0.04 mg/ml trypsin inhibitor, T6522). Cells were then filtered through a 40-mm Corning Cell Strainer. Filtered cells were gently shaken for 15 to 30 minutes before the experiment was carried out on Qube 384 using single-hole QChips (Sophion Biosciences). External solution contained 145 mM NaCl; 4 mM KCl; 10 mM HEPES; 6mM CaCl2; 1 mM MgCl2; 10 mM Glucose; pH was adjusted to 7.4 with NaOH, and osmolarity was adjusted to approximately 305 mOsm. Internal solution contains (in mM): 120 mM KF; 20 mM KCl; 10 mM HEPES; 10 mM EGTA; pH was adjusted to 7.2 with KOH, and osmolarity was adjusted to approximately 300 mOsm. Experiments were carried out at room temperature. Voltage-dependent sodium currents were evoked by 100 ms voltage steps to −10 mV from a holding potential of −100 mV. Steps were applied every 10 seconds. After recording control currents for approximately 1.5 minutes, 10 nM Protoxin II was applied and current was recorded in the presence of Protoxin II for 2 minutes. After that, 1 μM TTX, in the continuing presence of Protoxin II, was added and current was recorded for another 2 minutes in the presence of TTX and Protoxin II. Effects of Protoxin and TTX on peak sodium current were quantified in cells that met acceptance criteria of control sodium current of at least 150 pA in cells with a capacitance of at least 4 pF and total resistance (parallel combination of cell input and seal resistances) greater than 0.5 GOhm.

### Calcium imaging

Calcium imaging was performed using the FLIPR Tetra instrument and FLIPR Calcium 6 Assay Kit (Molecular Devices) according to the manufacturer’s instructions. In brief, Day 14 nociceptors were plated at a density of 2 x 10^5^ cells/cm^2^ on Geltrex coated black 96-well or 384-well plates and matured for 14 days as described above. Baseline signals were recorded for 2 minutes and then 40 mM KCL, 10 μM α,β-me-ATP, 1 μM capsaicin, 100 μM mustard oil or 250 μM menthol was applied and calcium signals were recorded every 1 s for 4 minutes.

### RNAscope

RNAscope experiments were performed by Advanced Cell Diagnostics according to the protocol for fixed cells using the RNAscope Fluorescent Multiplex Reagent kit v2 assay (Advanced Cell Diagnostics). In brief, 6-week nociceptors cultured on 4-well chamber slides were fixed by 10% Neutral Buffered Formalin at room temperature for 30 min and then sequentially dehydrated by 50%, 70% and 100% ethanol. Samples were then washed with 1X PBS and treated with hydrogen peroxide (RNAscope, ref 322335) for 10 min at room temperature and washed in autoclaved distilled water. Using a steamer, samples were treated with distilled H_2_O for 10 s at 99°C and then moved to RNAscope 1X target Retrieval Reagent (RNAscope, ref 322000) for 5 min at 99°C. Samples were washed with autoclaved distilled water and transferred to 100% ethanol for 3 min. Then, samples were treated with RNAscope Protease III (RNAscope, ref 322337) for 30 min at 40°C and washed with autoclaved distilled water. Samples were treated with gene specific probes and negative (ref 320871) and positive (ref 320881) controls and hybridized for 2 h at 40°C in RNAscope HybEZ oven (Advanced Cell Diagnostics). A series of incubations was then performed to amplify the signal of the hybridized probe and label target probes for the assigned fluorescence detection channel (target probe was labeled for the assigned fluorescence detection channels Opal 520, Opal 570, Opal 650 and Opal 780, PerkinElmer). Nuclei were stained using a DAPI nuclear stain (RNAscope ref 323108) for 30 s at room temperature. The samples were finally mounted with ProLong Gold Antifade Mountant (ref P36934).

### HotSpot kinase inhibitor profiling

Human kinase profiling against a panel of 374 wild-type kinases was performed by by Reaction Biology (Malvern, PA). Determination of IC_50_ of CHIR99021, CHIR98014, SU5402, and PD173074 and other experimental details are presented in Supplementary Table 1.

### FAAH inhibition assay

Nociceptors (day 28, WA09) cultured in black 96-well plates were washed 3 times in PBS and 250 μl PBS was added into each well. Twenty-one known FAAH inhibitors were added to the nociceptors to final concentration of 10 μM (8 replicates for each compound). Nociceptors were incubated at 37 °C for 30 min. Then, FAAH substrate AAMCA was added to a final concentration of 100 μM. Fluorescence signals were measured at 40 min after administration of AAMCA using a fluorescent plate reader (355 nm excitation, 460 nm emission).

### Bioinformatics analysis

Bioinformatics analysis was carried out using the computational resources of the NIH HPC Biowulf cluster (http://hpc.nih.gov) primarily using R language 4.0.3 (https://cran.r-project.org/).

For R scripts and program parameters, see https://github.com/cemalley/Deng_methods. Bulk RNA-Seq samples were quality trimmed using Trimmomatic 0.36 and the TruSeq3 paired-end adapters. STAR aligner 2.7.6a followed by HTSeq-count 0.11.4 produced deduplicated counts of reads in genes^47,48^. Counts were imported into DESeq2 1.30.0^49^. UCSC gene counts were normalized using the median-of-ratios method. Samples from two separate libraries were merged with batch correction using RUVSeq 1.24.0 RUVg with a set of housekeeping genes to adjust for systematic library size differences^50,51^. This produced a batch correction variable in the model for differential expression (DE) testing. The method is analogous to ERCC spike-ins batch correction. DRG samples from the SRA were also processed with batch correction when merged with nociceptors. DE tests used the conservative lfcShrink adjustment and negative binomial distribution. Principal Component Analysis used base prcomp. Heatmaps were made using scaled variance stabilizing transformed counts and ComplexHeatmap 2.6.2^52^.

Gene set enrichment tests were performed using Enrichr 3.0 and the ARCHS4 Tissues database^53^. Counts were visualized using the Integrative Genomics Browser (Extended Data Fig. 10g). For the four-way Venn diagram (Fig. 3c) package gplots 3.1.1 was used with a minimum threshold of 10 mean normalized counts across sample replicates to call a gene expressed in a sample.

Ion channel gene lists were taken from the HUGO Gene Nomenclature Committee hierarchy (https://www.genenames.org/data/genegroup/#!/group/177) with a threshold of 100 mean normalized counts.

### Data and code availability

RNA-Seq samples have been deposited to the Sequence Read Archive under PRJNA783035. Data analysis scripts are available at https://github.com/cemalley/Deng_methods. Data representing human dorsal root ganglia was accessed through dbGaP (phs001158.v2.p1)

