## Supplementary figures and images for "Scalable Generation of Pseudo-Unipolar Sensory Neurons from Human Pluripotent Stem Cells"

### Ext Data Fig 1

#### Extended Data Figure 1 (Deng et al.)

**a**

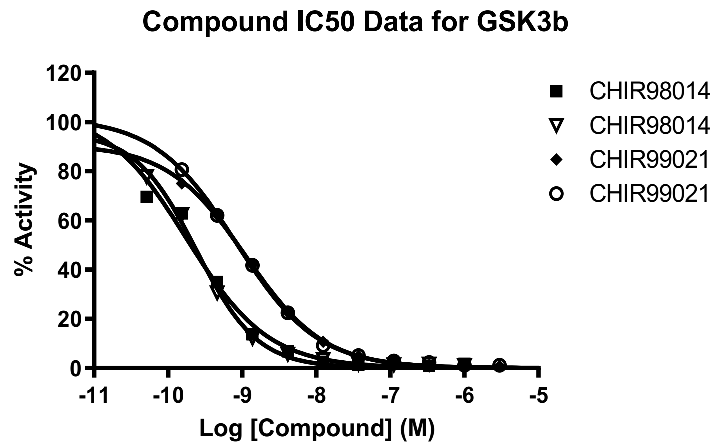


| | CHIR98014 | CHIR98014 | CHIR99021 | CHIR99021 |
|---|---|---|---|---|
| HillSlope | -0.7856 | -1.051 | -0.8216 | -0.7889 |
| EC50 | 1.784e-010 | 2.398e-010 | 1.120e-009 | 8.280e-010 |

**b**

**CHIR99021 (1.5 µM)**

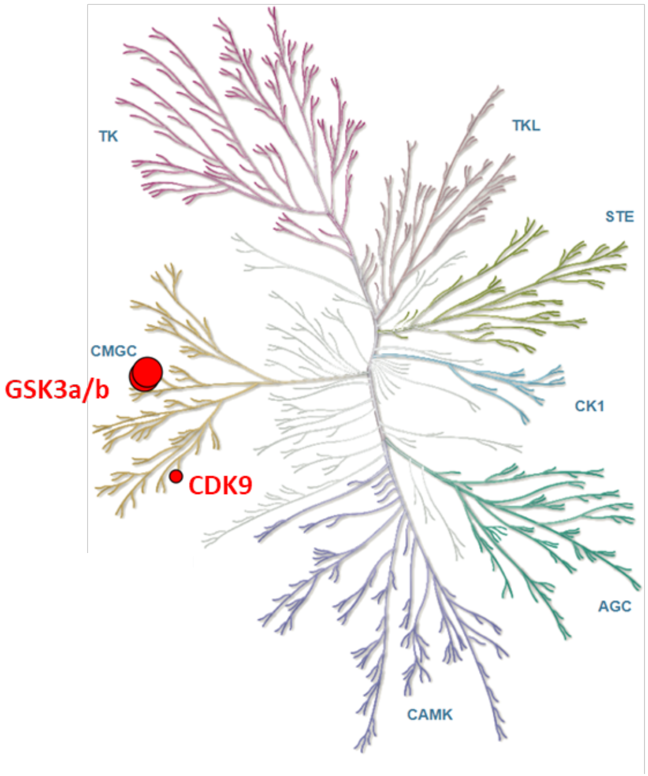


**CHIR98014 (0.5 µM)**

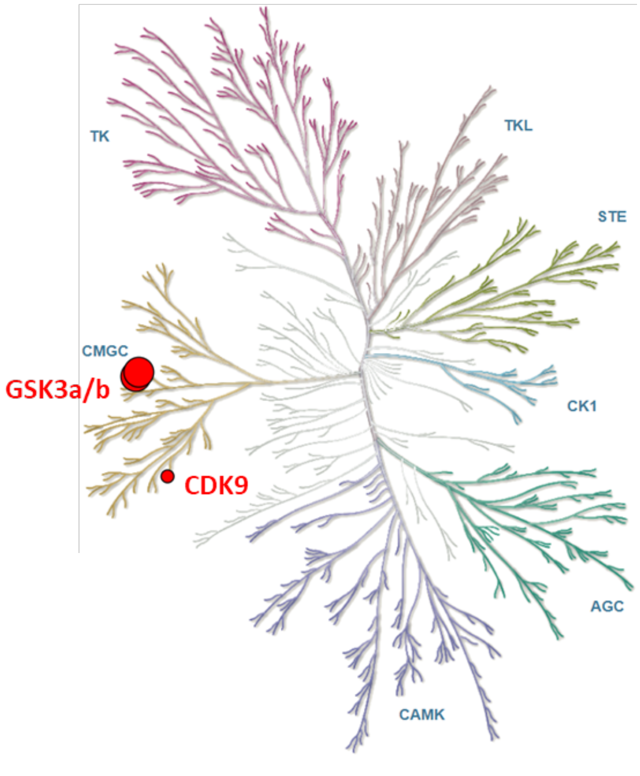

### Ext Data Fig 2

#### Extended Data Figure 2 (Deng et al.)

**a**

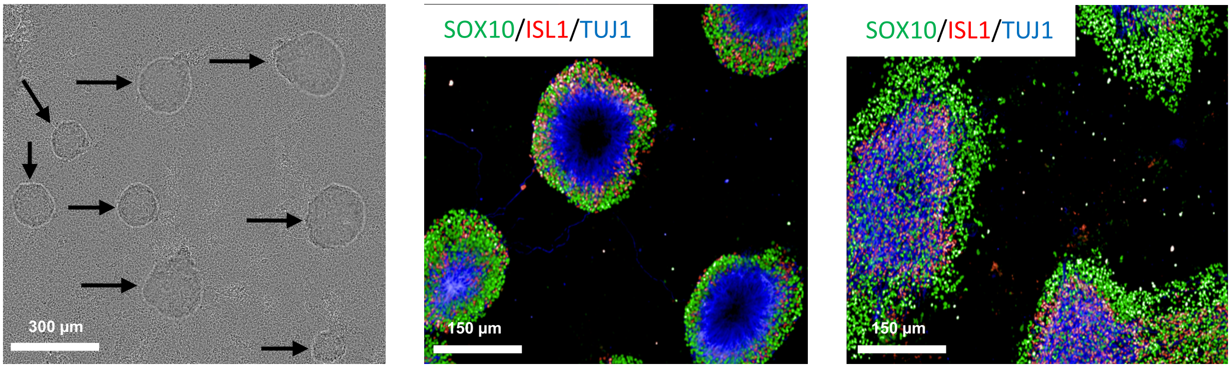


**b**

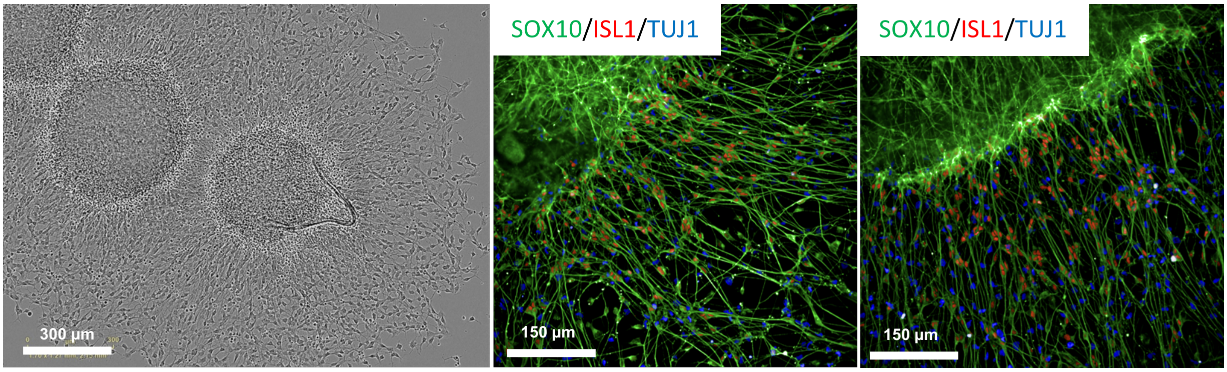


**c**

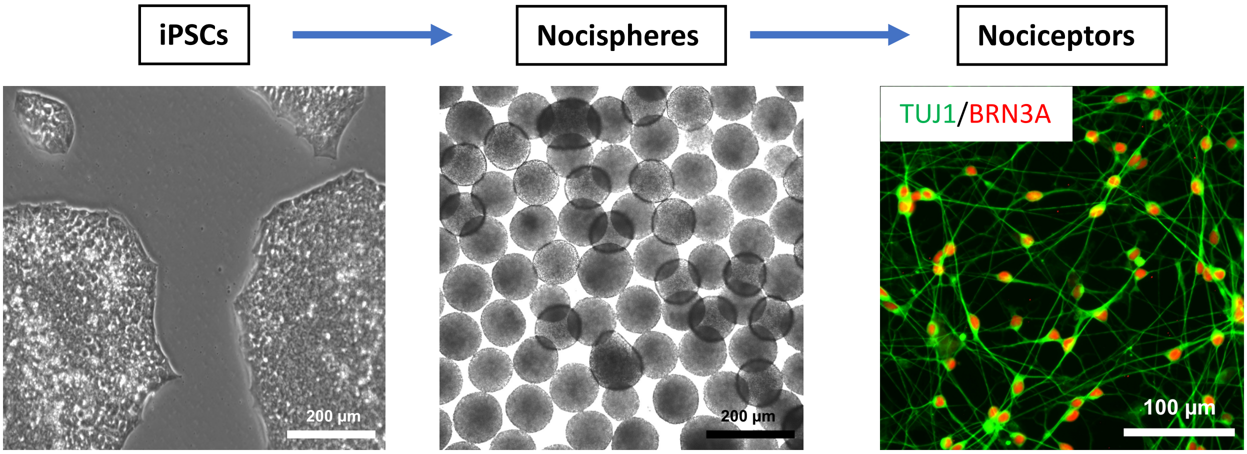

### Ext Data Fig 3

#### Extended Data Figure 3 (Deng et al.)

**a**

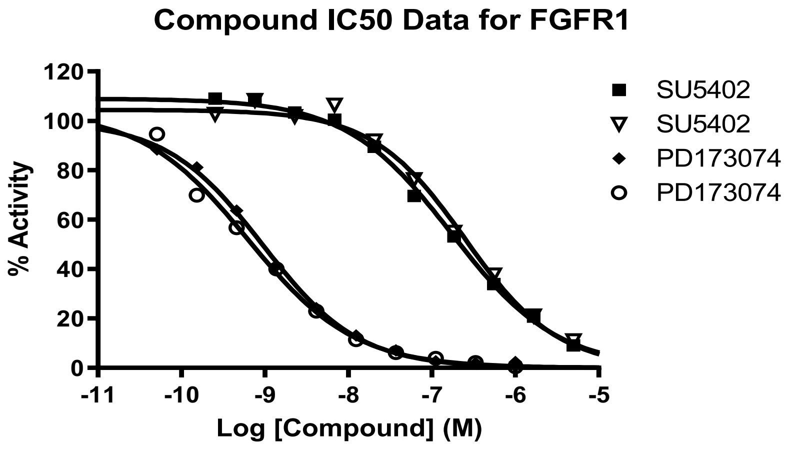


| | SU5402 | SU5402 | PD173074 | PD173074 |
|---|---|---|---|---|
| HillSlope | -0.6880 | -0.7803 | -0.7515 | -0.6814 |
| EC50 | 1.715e-007 | 2.501e-007 | 9.346e-010 | 6.293e-010 |

**b**

**SU5402 (0.5 µM)**

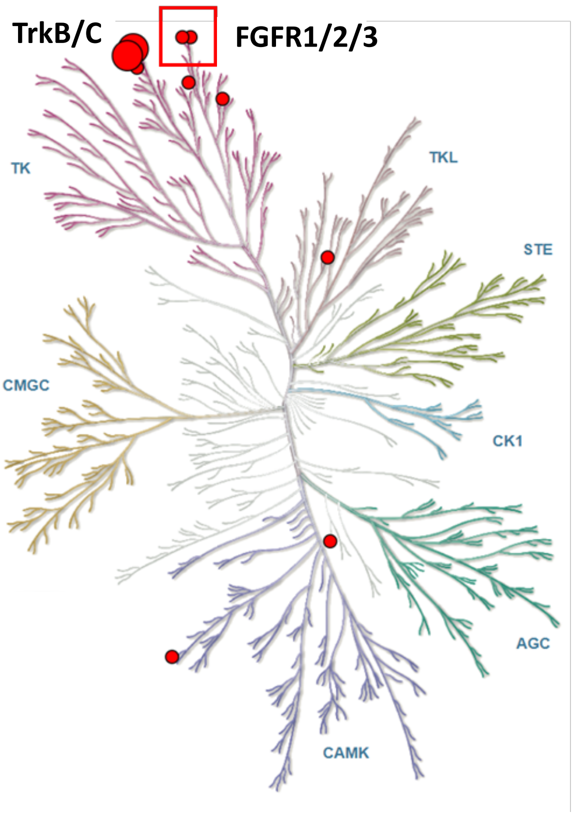


**PD173074 (25 nM)**

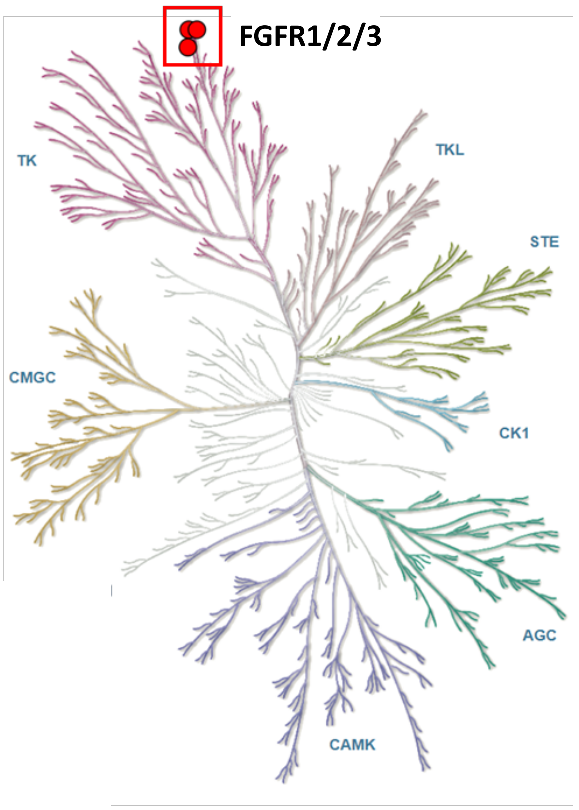

### Ext Data Fig 4

Extended Data Figure 4 (Deng et al.)

a

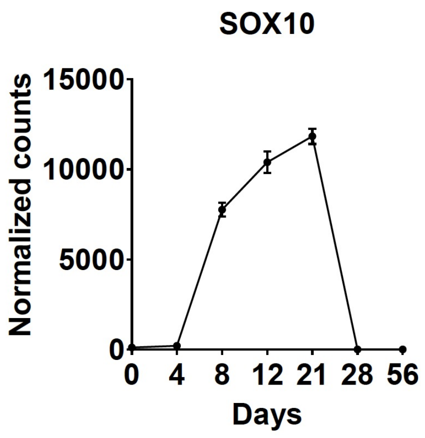


b

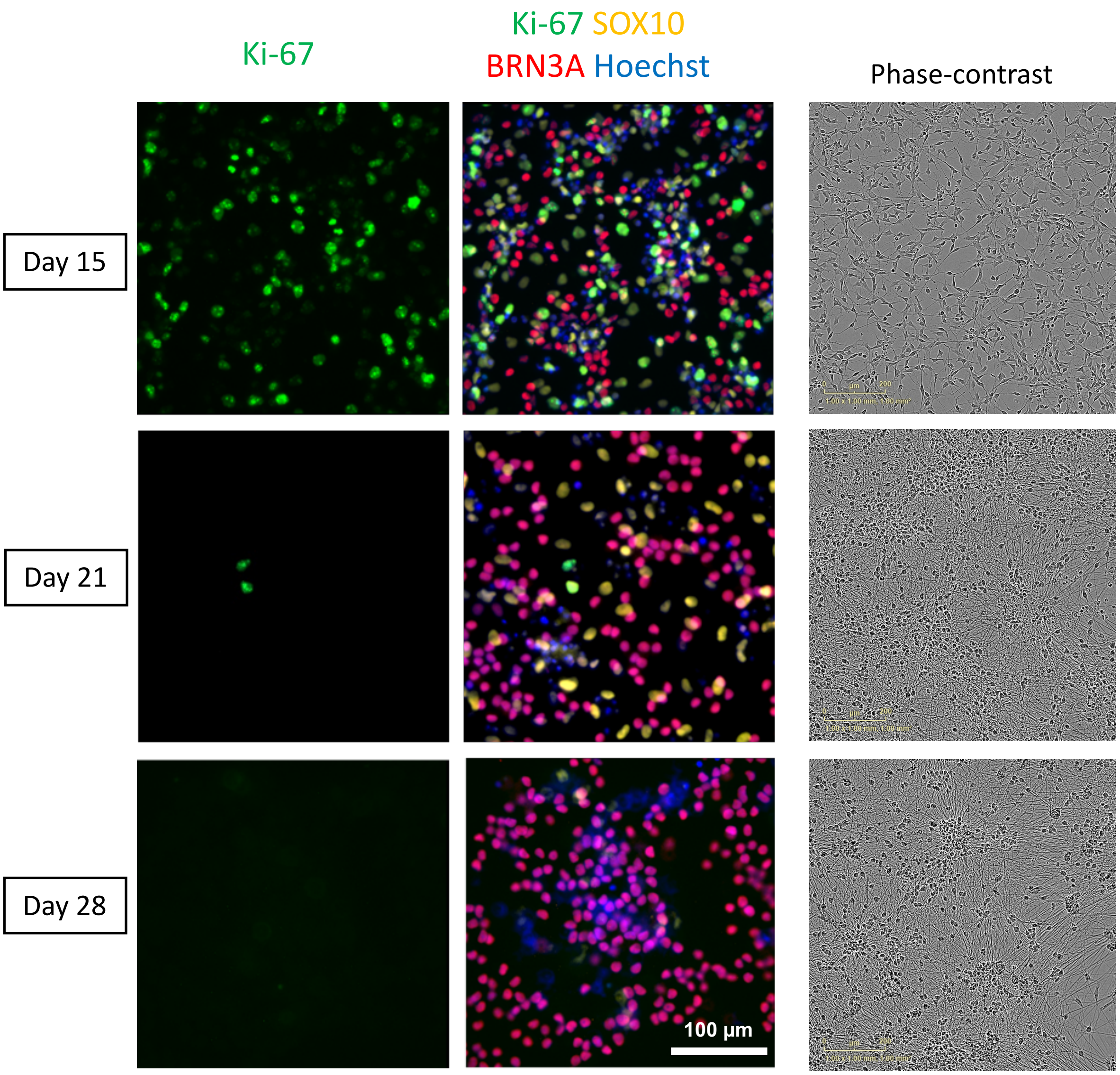

### Ext Data Fig 6

#### Extended Data Figure 6 (Deng et al.)

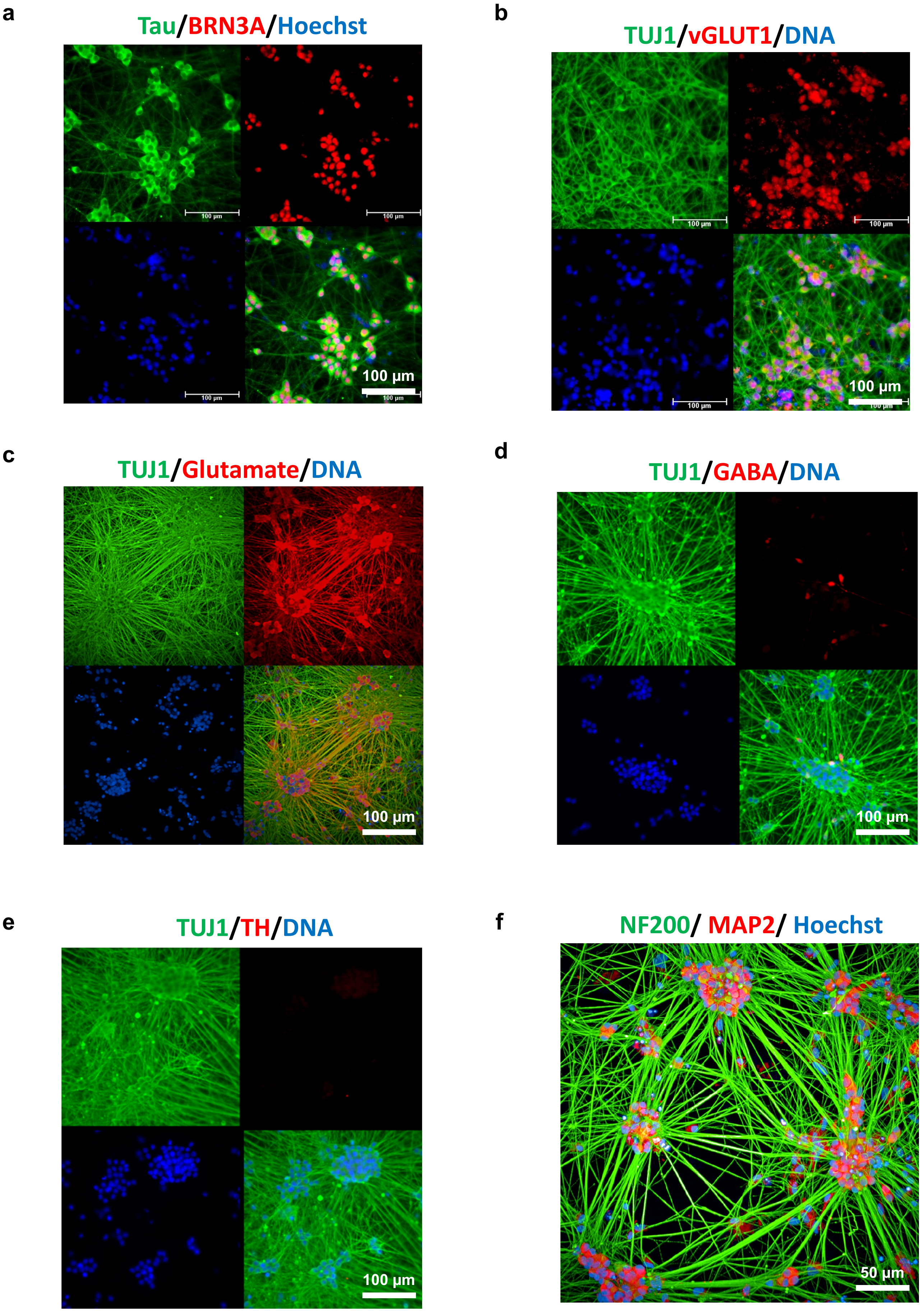

### Ext Data Fig 7

#### Extended Data Figure 7 (Deng et al.)

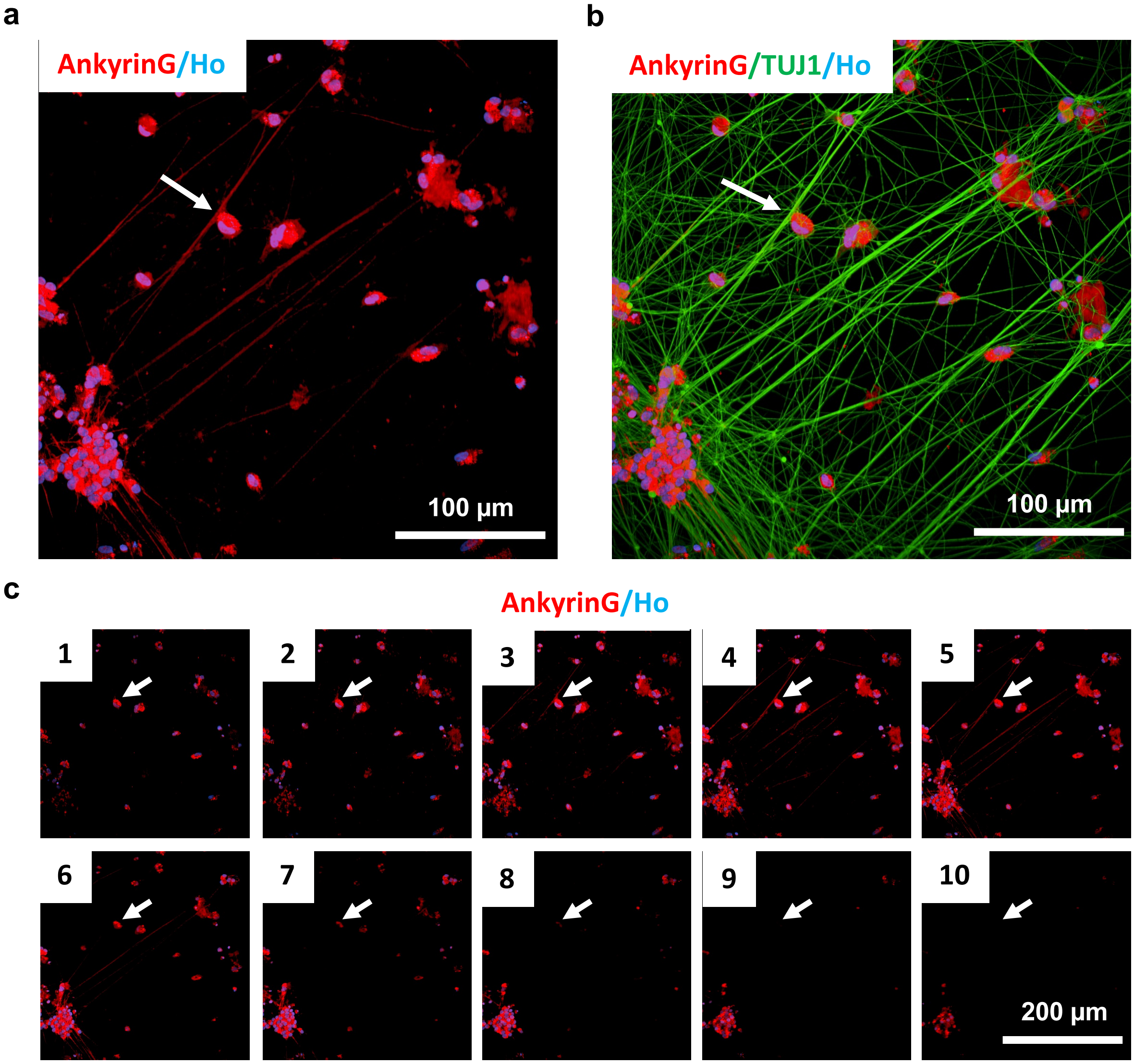

### Ext Data Fig 8

#### Extended Data Figure 8 (Deng et al.)

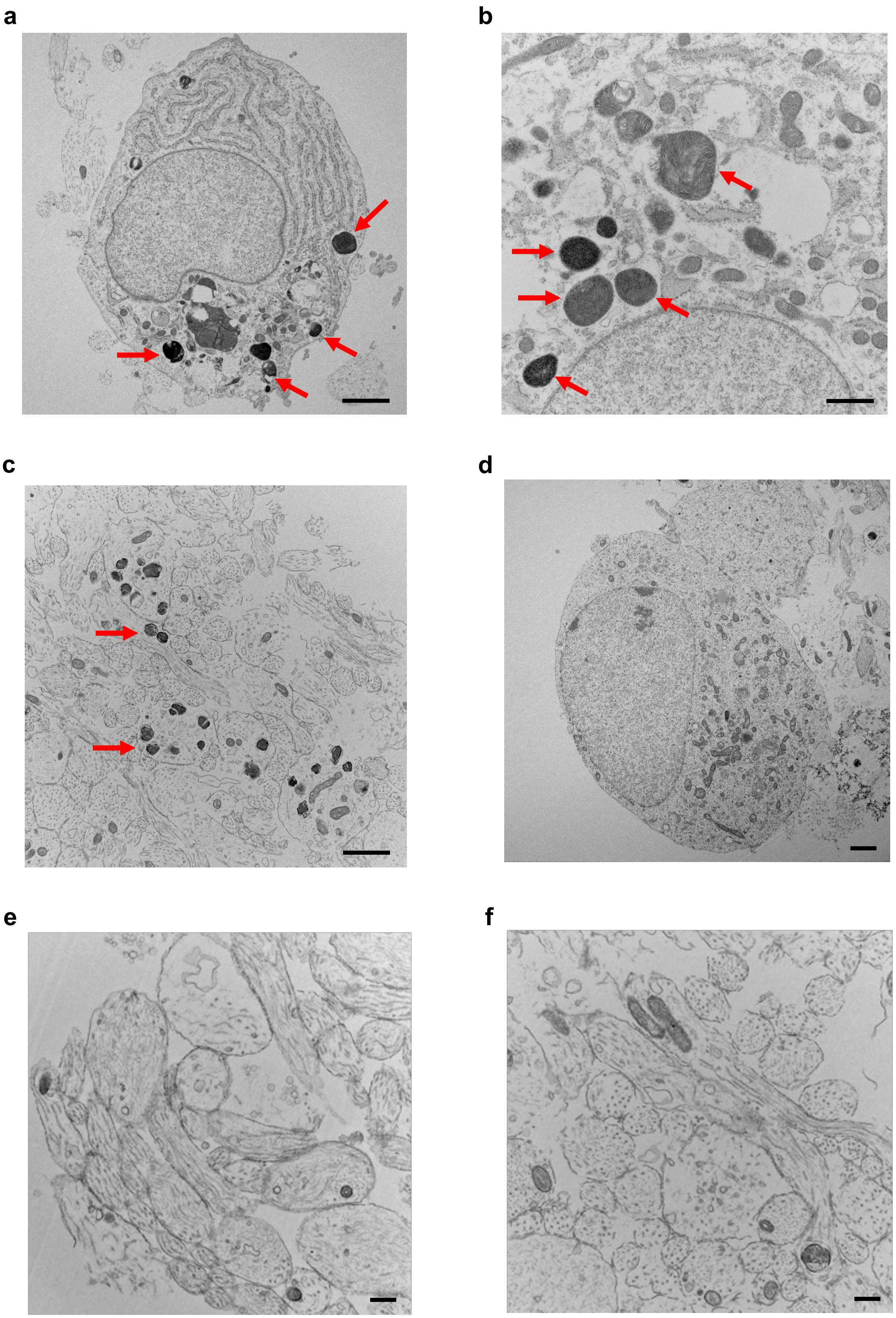

### Ext Data Fig 9

#### Extended Data Figure 9 (Deng et al.)

**a**

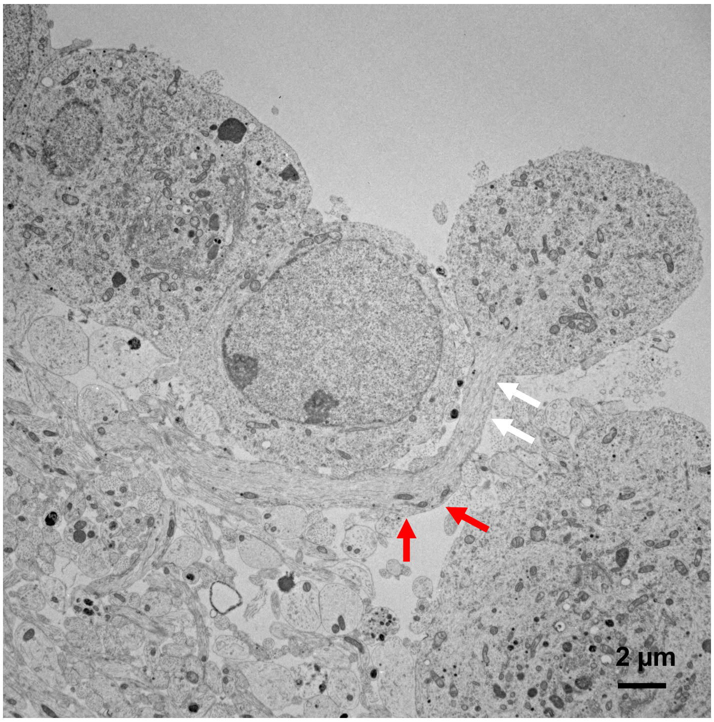


**b**

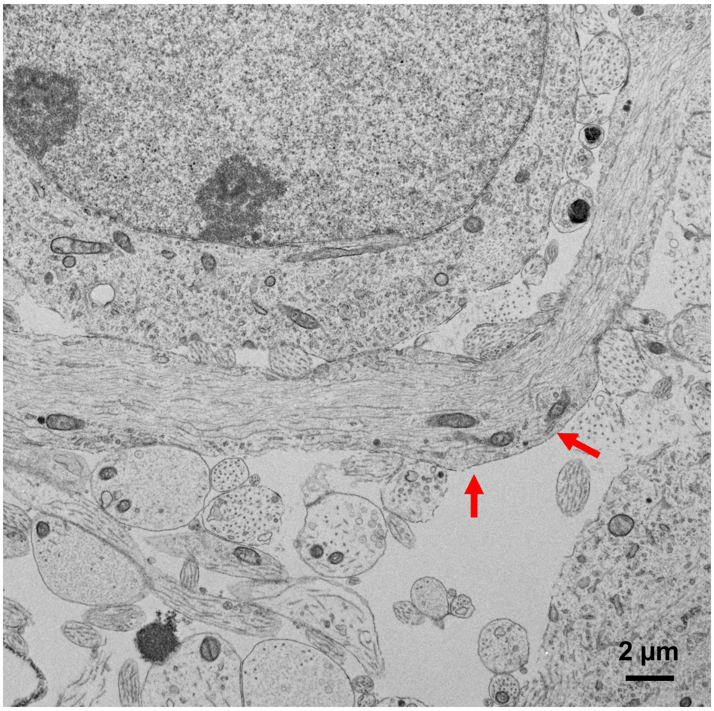

### Ext Data Fig 10

#### Extended Data Figure 10 (Deng et al.)

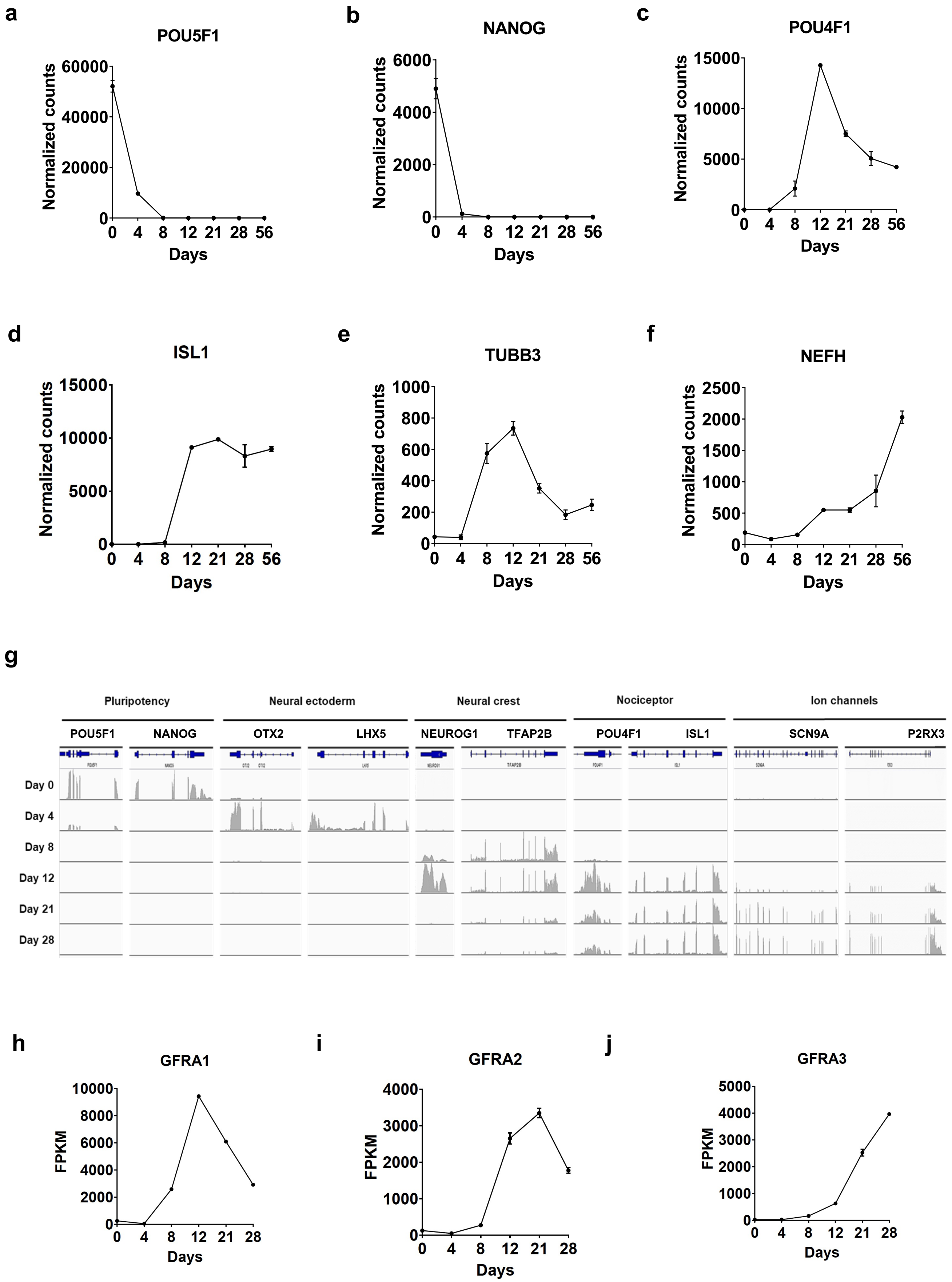

### Ext Data Fig 11

#### Extended Data Figure 11 (Deng et al.)

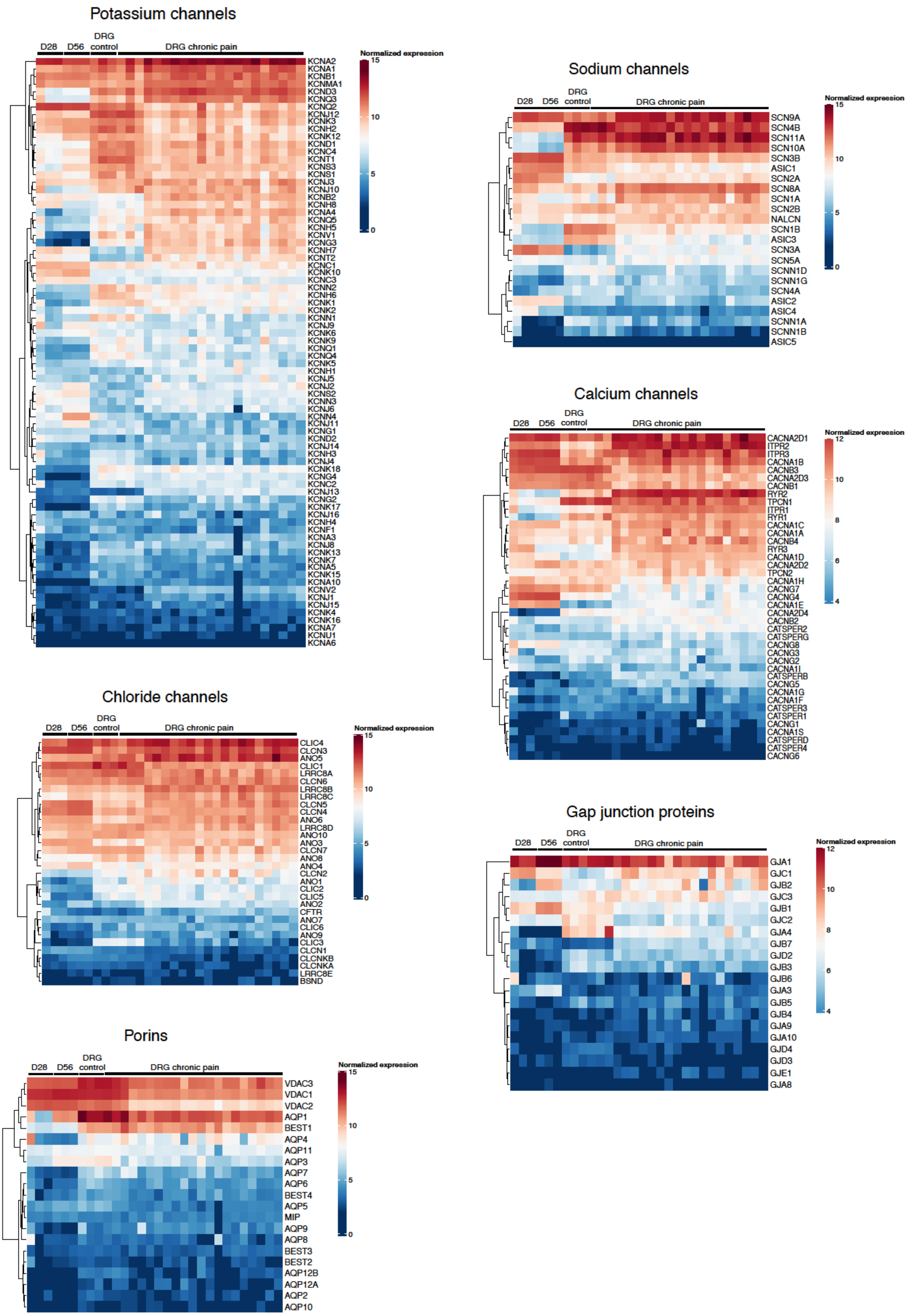

### Ext Data Fig 12

#### Extended Data Figure 12 (Deng et al.)

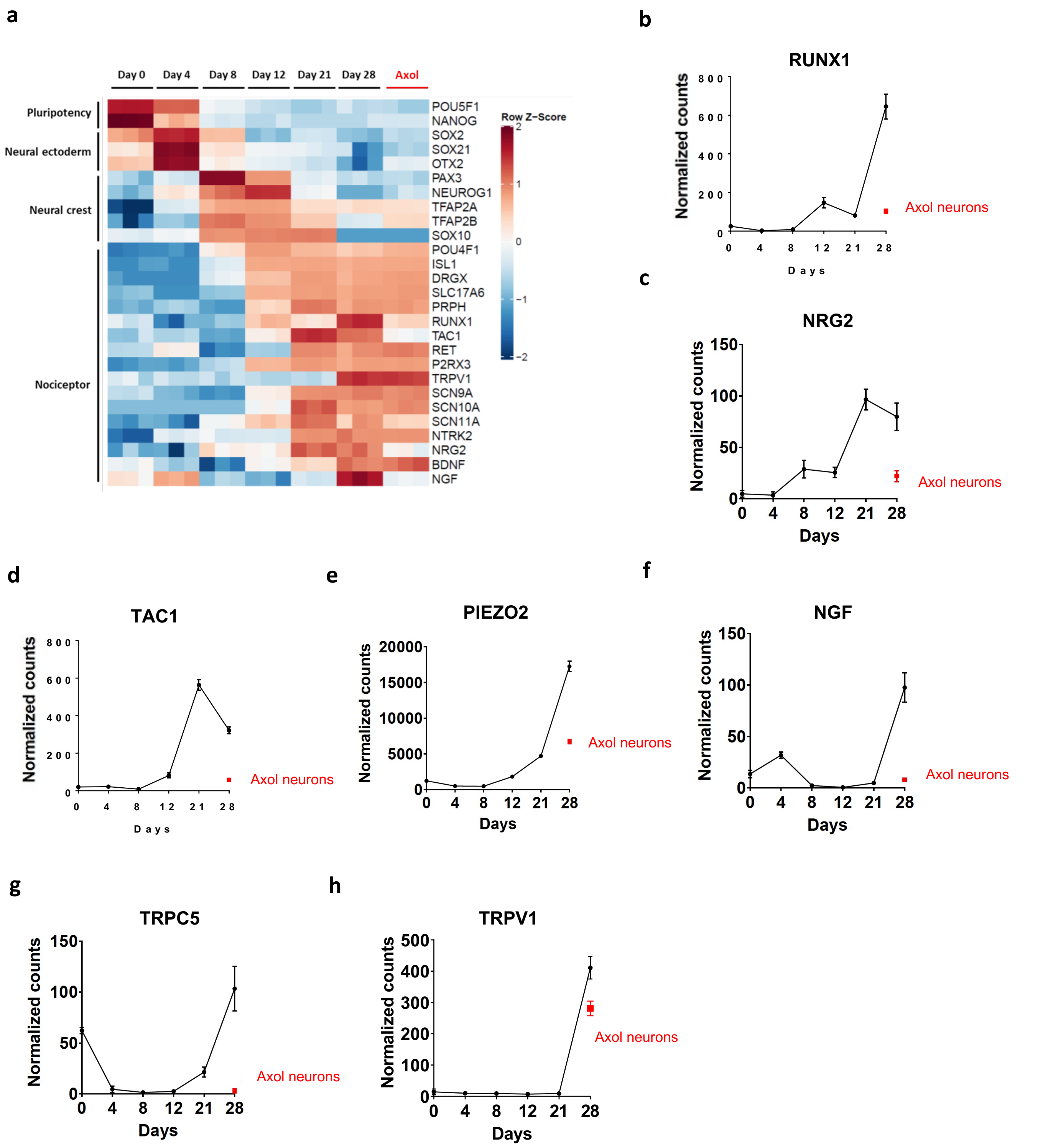

### Ext Data Fig 13

#### Extended Data Figure 13 (Deng et al.)

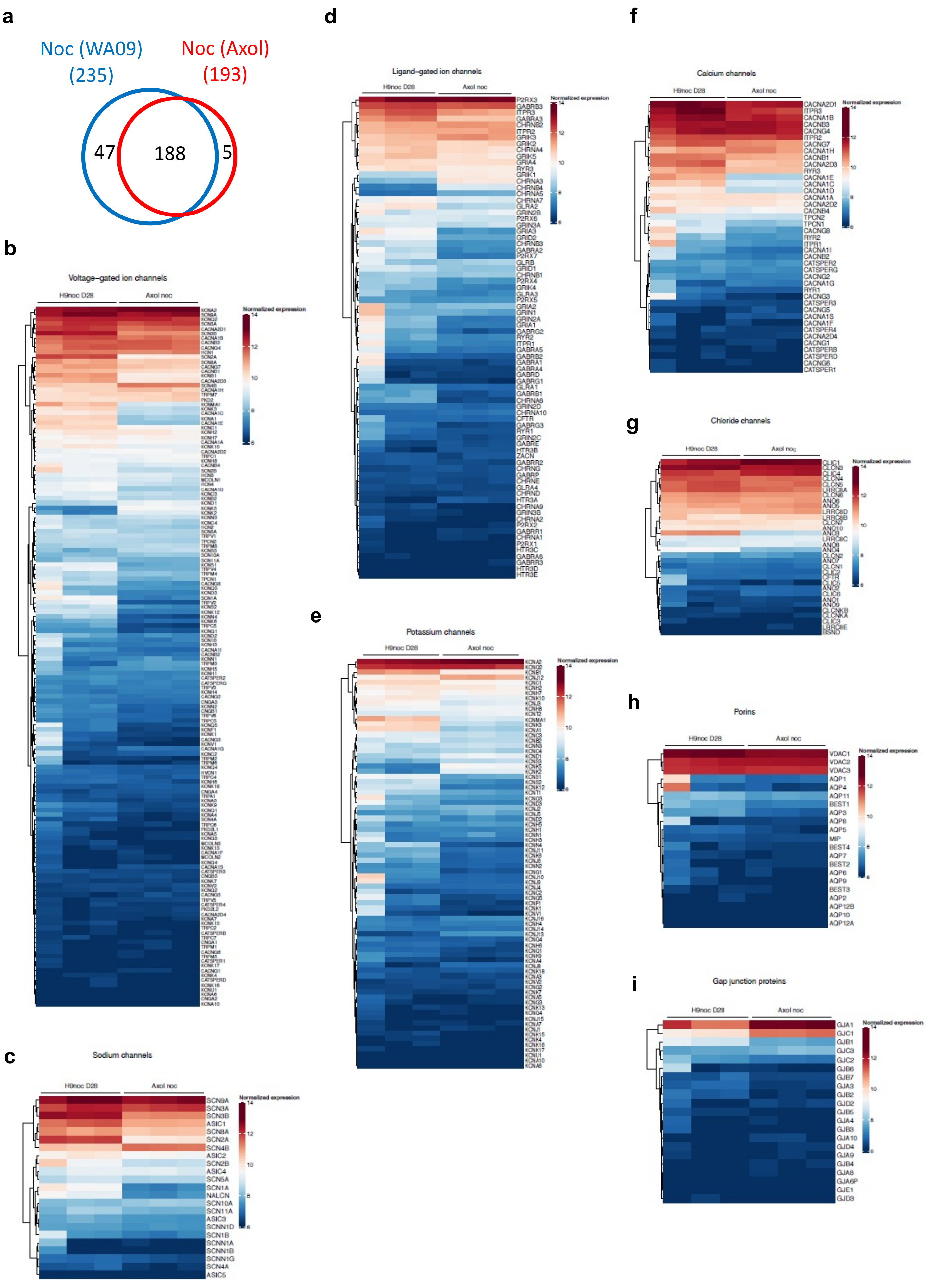
