## Supplementary material for "Scalable Generation of Pseudo-Unipolar Sensory Neurons from Human Pluripotent Stem Cells": Ext Data Fig 5

### Extended Data Figure 5 (Deng et al.)

**a**

CompacT SelecT™

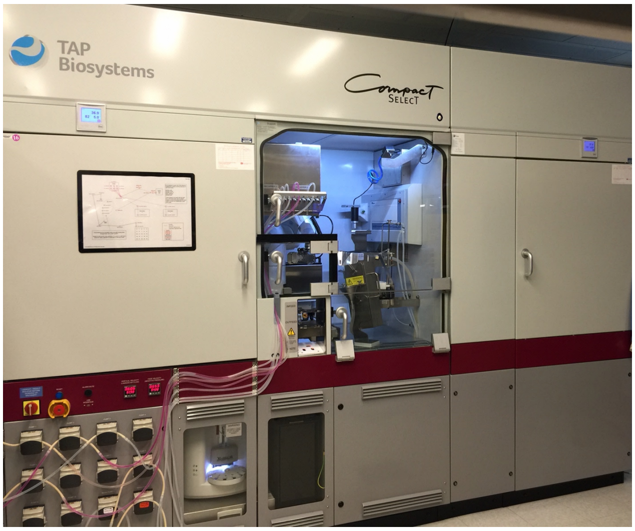


**b**

**2.5 x $10^7$ iPSCs per flask**

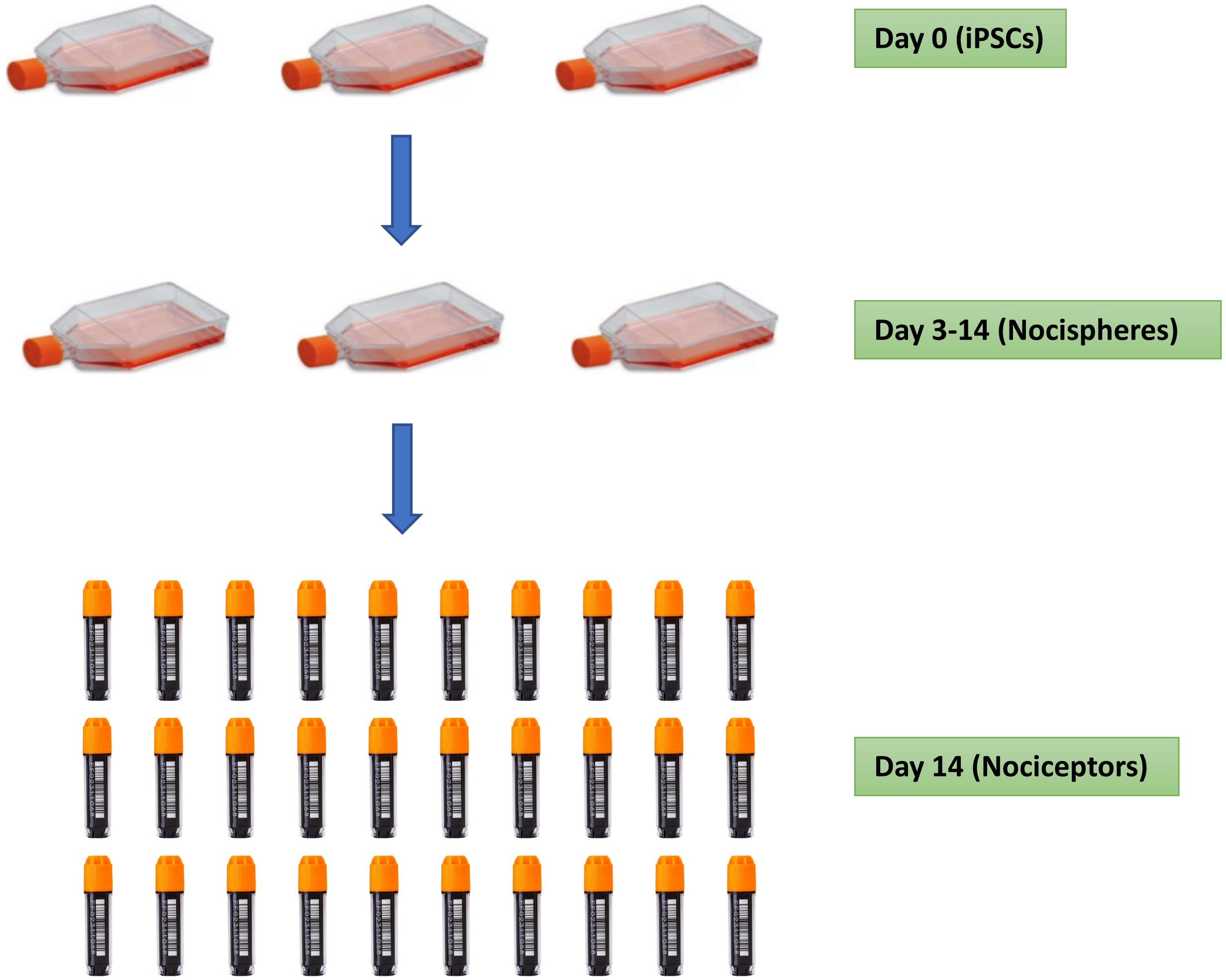


**30 vials, 5 x $10^6$ nociceptors per vial (1.5 x $10^8$ nociceptors in total)**
