## Supplementary material for "Scalable Generation of Pseudo-Unipolar Sensory Neurons from Human Pluripotent Stem Cells": Ext Data Fig 14

### Extended Data Figure 14 (Deng et al.)

**a**

Nociceptors RNA-seq normalized counts (3 replicates at each time point)

| Gene | Day 0 - 1 | Day 0 - 2 | Day 0 - 3 | Day 4 -1 | Day 4 -2 | Day 4 -3 | Day 8 - 1 | Day 8 - 2 | Day 8 - 3 | Day 12 - 1 | Day 12 - 2 | Day 12 - 3 | Day 21 - 1 | Day 21 - 2 | Day 21 - 3 | Day 28 -1 | Day 28 -2 | Day 28 -3 |
|---|---|---|---|---|---|---|---|---|---|---|---|---|---|---|---|---|---|---|
| SCN1A | 32.63378 | 21.20595 | 35.01409 | 10.56849 | 4.994754 | 2.337063 | 453.6262 | 346.2216 | 354.2262 | 67.31298 | 70.97562 | 80.17315 | 241.2549 | 227.647 | 209.3765 | 271.9445 | 272.5455 | 242.3641 |
| SCN1B | 62.75726 | 54.94269 | 77.27247 | 248.3594 | 292.1931 | 261.7511 | 22.84448 | 28.76302 | 36.40869 | 227.7683 | 220.813 | 251.6204 | 49.03556 | 55.2213 | 57.55372 | 45.06945 | 49.17488 | 35.18189 |
| SCN2A | 37.65436 | 18.31423 | 18.11073 | 35.66864 | 47.45016 | 42.06714 | 42.42547 | 49.00367 | 42.4768 | 196.46 | 219.8273 | 187.4818 | 1824.123 | 1718.622 | 1839.735 | 2718.681 | 2588.766 | 2688.483 |
| SCN2B | 2.510291 | 1.927814 | 2.414765 | 3.963183 | 3.746066 | 4.674126 | 22.84448 | 28.76302 | 27.30652 | 47.74525 | 43.37399 | 43.17016 | 330.4996 | 303.1536 | 356.2377 | 535.4862 | 451.7421 | 496.4556 |
| SCN3A | 2.510291 | 0 | 0 | 29.06334 | 21.22771 | 19.86504 | 181.668 | 262.0631 | 304.1642 | 4304.9 | 4489.208 | 4294.814 | 5125.196 | 5465.781 | 5492.411 | 4810.973 | 4591.6 | 4753.465 |
| SCN3B | 15.06174 | 9.639069 | 15.69597 | 26.42122 | 37.46066 | 28.04476 | 60.91862 | 117.1827 | 124.7756 | 1379.916 | 1437.256 | 1337.042 | 3363.839 | 3394.419 | 3393.685 | 4678.82 | 4417.404 | 4490.578 |
| SCN4A | 340.1444 | 350.8621 | 441.9019 | 1052.886 | 991.4587 | 983.9036 | 50.0403 | 49.00367 | 46.26937 | 359.2634 | 363.7501 | 397.1655 | 14.71067 | 23.66627 | 20.83842 | 6.875001 | 5.834308 | 4.886374 |
| SCN4B | 85.34988 | 80.00427 | 94.17582 | 352.7233 | 375.8553 | 427.6825 | 149.0331 | 116.1174 | 137.2911 | 236.3781 | 181.3821 | 249.1535 | 335.4032 | 397.8187 | 333.4147 | 100.8333 | 90.84851 | 115.3184 |
| SCN5A | 520.8853 | 515.6902 | 620.5945 | 368.576 | 353.3789 | 338.8742 | 233.884 | 232.2348 | 207.0744 | 335.7822 | 283.9025 | 319.4592 | 559.0053 | 546.5781 | 505.0835 | 600.4167 | 488.4149 | 544.3421 |
| SCN7A | 6.275726 | 14.4586 | 4.829529 | 9.247426 | 9.989508 | 1.168532 | 154.4722 | 134.2274 | 107.709 | 109.5793 | 119.2785 | 123.3433 | 743.379 | 703.2263 | 798.806 | 2245.07 | 2116.187 | 2130.459 |
| SCN8A | 1541.318 | 1461.283 | 1281.033 | 1034.391 | 1125.068 | 1155.678 | 4882.192 | 5517.174 | 5327.046 | 2006.866 | 2018.862 | 2054.9 | 1504.411 | 1559.72 | 1598.604 | 1203.125 | 1186.031 | 1161.002 |
| SCN9A | 138.066 | 125.3079 | 173.8631 | 48.87925 | 44.95279 | 28.04476 | 14.14182 | 20.24065 | 15.9288 | 733.3984 | 769.8883 | 873.2707 | 6191.229 | 6405.67 | 6840.955 | 5354.098 | 5062.512 | 5152.193 |
| SCN10A | 0 | 0 | 0 | 0 | 0 | 0 | 0 | 0 | 0 | 14.87147 | 23.65854 | 14.8012 | 478.587 | 367.3907 | 470.3528 | 263.5417 | 254.2091 | 258.9778 |
| SCN11A | 20.08232 | 16.38642 | 25.35503 | 23.7791 | 16.23295 | 8.179721 | 75.06044 | 75.6361 | 87.98766 | 152.6283 | 170.5386 | 149.2454 | 323.6347 | 300.8997 | 296.6994 | 221.5278 | 175.0292 | 241.3869 |
| KCNQ1 | 150.6174 | 170.6115 | 155.7523 | 60.7688 | 62.43443 | 57.25805 | 32.63498 | 49.00367 | 47.40714 | 21.91585 | 13.80082 | 18.5015 | 11.76853 | 7.888757 | 10.91536 | 17.56945 | 4.167363 | 2.931824 |
| KCNQ2 | 1479.816 | 1620.327 | 1668.602 | 697.5202 | 508.2162 | 621.6588 | 363.3361 | 382.4417 | 439.9383 | 1753.268 | 1519.075 | 1583.728 | 1788.817 | 1871.889 | 1591.658 | 2078.542 | 1970.329 | 1857.799 |
| KCNQ3 | 84.09473 | 98.3185 | 72.44294 | 11.88955 | 16.23295 | 12.85385 | 18.49315 | 35.15481 | 29.58206 | 177.675 | 170.5386 | 155.4126 | 203.0072 | 205.1077 | 246.0918 | 241.3889 | 267.5447 | 267.7733 |
| KCNQ4 | 184.5064 | 169.6476 | 150.9228 | 165.1326 | 197.2928 | 181.1224 | 58.74296 | 47.93837 | 43.99383 | 36.78732 | 36.47358 | 39.46986 | 37.26702 | 32.68199 | 32.74608 | 47.36112 | 46.67446 | 63.52286 |
| KCNQ5 | 72.79843 | 57.83441 | 67.61341 | 1.321061 | 1.248689 | 0 | 0 | 3.195891 | 3.413314 | 2.348127 | 2.957318 | 3.700299 | 1.961422 | 9.015722 | 16.8692 | 42.77778 | 32.50543 | 37.13644 |
| FGF11 | 45.18523 | 56.87051 | 59.16173 | 113.6112 | 98.6464 | 113.3476 | 13.05399 | 3.195891 | 9.102172 | 3.913545 | 12.81504 | 8.634032 | 14.71067 | 15.77751 | 5.953834 | 29.79167 | 26.67112 | 23.4546 |
| FGF12 | 646.3998 | 624.6117 | 748.577 | 177.0222 | 201.0389 | 190.4706 | 886.5835 | 609.35 | 585.9523 | 238.7263 | 267.1444 | 240.5195 | 203.9879 | 178.0605 | 220.2918 | 198.6111 | 163.3606 | 148.5458 |
| FGF13 | 2190.228 | 2151.44 | 2281.953 | 397.6393 | 418.3107 | 413.6602 | 1165.069 | 1232.549 | 1142.323 | 4722.866 | 4881.546 | 4904.13 | 22265.08 | 22277.85 | 22905.39 | 26091.39 | 26961.17 | 26017.99 |
| FGF14 | 5.020581 | 3.855628 | 4.829529 | 6.605304 | 12.48689 | 9.348252 | 10.87833 | 18.11005 | 14.41177 | 18.78502 | 22.67277 | 23.43523 | 33.34418 | 36.06289 | 46.63836 | 110.7639 | 128.3548 | 126.0684 |

**b**

Nociceptors RNA-seq normalized counts (3 replicates at each time point)

| Gene | Day 0 - 1 | Day 0 - 2 | Day 0 - 3 | Day 4 -1 | Day 4 -2 | Day 4 -3 | Day 8 - 1 | Day 8 - 2 | Day 8 - 3 | Day 12 - 1 | Day 12 - 2 | Day 12 - 3 | Day 21 - 1 | Day 21 - 2 | Day 21 - 3 | Day 28 -1 | Day 28 -2 | Day 28 -3 |
|---|---|---|---|---|---|---|---|---|---|---|---|---|---|---|---|---|---|---|
| PDGFA | 2737.472 | 2787.619 | 2486 | 2778.191 | 2863.243 | 2807.981 | 10.87833 | 24.50183 | 13.274 | 75.14007 | 74.91871 | 67.83882 | 134.3574 | 114.9505 | 141.8997 | 119.9306 | 123.3539 | 96.7502 |
| PDGFB | 1501.154 | 1541.287 | 926.0622 | 22.45804 | 37.46066 | 28.04476 | 1.087833 | 2.130594 | 2.6548 | 3.913545 | 3.94309 | 11.1009 | 52.9584 | 52.96737 | 62.51525 | 165 | 189.1983 | 171.0231 |
| PDGFC | 121.7491 | 118.5605 | 124.3604 | 237.791 | 209.7797 | 252.4028 | 733.1991 | 776.6016 | 796.8193 | 375.7003 | 410.0814 | 389.7649 | 404.053 | 382.0412 | 440.5837 | 888.4029 | 980.1637 | 913.7519 |
| PDGFD | 160.6586 | 194.7092 | 195.5959 | 88.51108 | 98.6464 | 75.95455 | 143.5939 | 157.664 | 178.2509 | 104.883 | 108.435 | 94.97435 | 240.2742 | 210.7425 | 235.1764 | 417.0834 | 415.0693 | 444.66 |
| PDGFRA | 16.31689 | 10.60298 | 24.14765 | 29.06334 | 26.22246 | 23.37063 | 158.8235 | 151.2722 | 132.3607 | 82.96716 | 103.5061 | 82.64002 | 79.4376 | 101.4269 | 106.1767 | 728.7501 | 855.9763 | 797.4562 |
| PDGFRB | 428.0045 | 440.5054 | 377.9107 | 224.5804 | 182.3085 | 206.8301 | 253.465 | 307.8709 | 272.3066 | 1790.056 | 1640.325 | 1758.876 | 747.3019 | 829.4464 | 716.4446 | 2149.584 | 2076.18 | 2130.459 |
| PDGFRL | 57.73668 | 63.61785 | 84.51676 | 50.20031 | 54.9423 | 52.58392 | 40.2498 | 41.54659 | 31.8576 | 1.565418 | 0 | 0 | 5.884267 | 6.761791 | 5.953834 | 42.01389 | 42.5071 | 40.06827 |
| EGF | 151.8726 | 127.2357 | 155.7523 | 130.785 | 163.5782 | 141.3923 | 31.54714 | 19.17535 | 17.44583 | 3.130836 | 0.985773 | 0 | 41.18987 | 34.93592 | 42.66914 | 20.625 | 39.17321 | 29.31824 |
| EGFR | 345.1649 | 344.1148 | 340.4818 | 130.785 | 117.3767 | 125.0329 | 2073.409 | 2542.864 | 2432.555 | 399.1816 | 335.1627 | 339.1941 | 588.4267 | 623.2118 | 639.0448 | 844.8612 | 921.8206 | 936.2293 |
| NTRK1 | 1.255145 | 6.747348 | 6.036912 | 9.247426 | 6.243443 | 10.51678 | 219.7422 | 509.212 | 500.6194 | 3738.218 | 3417.673 | 3553.521 | 1460.279 | 1480.832 | 1295.951 | 16.80556 | 13.33556 | 10.75002 |
| NTRK2 | 1.255145 | 0 | 0 | 232.5067 | 144.8479 | 217.3469 | 344.8429 | 133.1621 | 146.7725 | 165.1516 | 169.5529 | 154.1791 | 5209.537 | 5473.67 | 5263.189 | 6686.32 | 7132.024 | 6748.082 |
| NTRK3 | 434.2803 | 458.8197 | 361.0073 | 284.0281 | 242.2456 | 271.0993 | 358.9847 | 416.5312 | 442.2138 | 478.2352 | 424.868 | 558.7452 | 464.8571 | 413.5962 | 358.2223 | 337.6389 | 321.7204 | 333.2507 |
| NRG1 | 32.63378 | 56.87051 | 20.5255 | 29.06334 | 39.95803 | 35.05595 | 471.0315 | 523.0609 | 565.8517 | 3436.093 | 3401.901 | 3269.831 | 2638.113 | 2759.938 | 2610.756 | 2177.848 | 2038.674 | 2026.868 |
| NRG2 | 2.510291 | 8.675162 | 3.622147 | 3.963183 | 0 | 7.011189 | 39.16197 | 22.37124 | 26.548 | 25.8294 | 22.67277 | 33.30269 | 112.7818 | 92.41115 | 106.1767 | 111.5278 | 115.0192 | 99.68203 |
| NRG3 | 139.3211 | 140.7304 | 101.4201 | 38.31077 | 36.21197 | 29.21329 | 115.3102 | 140.6192 | 127.0511 | 947.0779 | 898.0388 | 906.5733 | 1245.503 | 1250.931 | 1276.105 | 773.0557 | 746.7914 | 754.4561 |
| NRG4 | 100.4116 | 114.7049 | 96.59059 | 101.7217 | 104.8898 | 112.179 | 125.1007 | 122.5092 | 121.7415 | 120.5372 | 124.2073 | 111.009 | 125.531 | 134.1089 | 159.7612 | 158.8889 | 165.861 | 142.6821 |
| VEGFA | 1212.47 | 1219.342 | 2121.371 | 2626.269 | 2335.048 | 2418.86 | 800.6447 | 442.0983 | 634.4972 | 619.9056 | 548.0895 | 573.5464 | 212.8143 | 236.6627 | 217.3149 | 362.0834 | 390.0651 | 373.319 |
| VEGFB | 322.5723 | 344.1148 | 399.6436 | 622.2197 | 585.6349 | 660.2203 | 41.33764 | 31.95891 | 61.43966 | 133.0605 | 84.77644 | 94.97435 | 246.1585 | 244.5515 | 223.2688 | 564.514 | 539.2567 | 501.342 |
| VEGFC | 30.12349 | 46.26753 | 41.051 | 9.247426 | 9.989508 | 19.86504 | 21.75665 | 19.17535 | 19.34211 | 0.782709 | 6.900408 | 1.233433 | 11.76853 | 11.26965 | 20.83842 | 87.84723 | 82.51378 | 75.25016 |
| VEGFD | 6.275726 | 9.639069 | 3.622147 | 3.963183 | 0 | 0 | 6.526995 | 9.587674 | 10.23994 | 54.00692 | 106.4634 | 117.1761 | 7.845689 | 12.39662 | 11.90767 | 12.98611 | 8.334725 | 13.68185 |
| FLT1 | 2506.525 | 2338.438 | 2492.037 | 426.7027 | 518.2057 | 473.2553 | 7.614828 | 7.45708 | 4.930343 | 6.261672 | 8.871953 | 6.167166 | 0.980711 | 0 | 3.969222 | 21.38889 | 32.50543 | 17.59095 |
| KDR | 7859.72 | 7475.098 | 8003.737 | 4964.547 | 5109.634 | 5368.234 | 20.66882 | 19.17535 | 14.41177 | 2.348127 | 1.971545 | 3.700299 | 87.28329 | 55.2213 | 112.1305 | 79.44446 | 102.5171 | 71.34106 |
| FLT4 | 448.0869 | 562.9216 | 578.3361 | 240.4331 | 196.0441 | 207.9986 | 125.1007 | 105.4644 | 137.6703 | 39.13545 | 35.48781 | 34.53613 | 44.132 | 46.20557 | 44.65375 | 23.68056 | 44.17404 | 29.31824 |
| BDNF | 66.5227 | 60.72613 | 82.102 | 38.31077 | 27.47115 | 33.88742 | 4.35133 | 14.91416 | 9.102172 | 87.66341 | 75.90448 | 88.80719 | 142.2031 | 127.3471 | 131.9766 | 185.625 | 213.369 | 174.9322 |
| NGF | 16.31689 | 15.42251 | 9.659059 | 33.02652 | 29.96853 | 36.22448 | 1.087833 | 3.195891 | 2.6548 | 0.782709 | 0.985773 | 0 | 5.884267 | 4.507861 | 4.961528 | 99.30557 | 88.34809 | 86.97746 |
